# Effect-oriented automated muscle path modeling via gradient-specified optimization using muscle surface mesh and moment arm

**DOI:** 10.64898/2026.04.15.718668

**Authors:** Ziyu Chen, Tingli Hu, Sami Haddadin, David W. Franklin

## Abstract

There is more to musculotendon path modeling than aligning a cable to reflect the geometric features of a musculotendon unit. From the perspective of simulation accuracy, the key is to replicate the length– and moment arm–joint angle relations of the target muscle. In this study, we propose an effect-oriented approach of automated path modeling, via hybrid calibration based on muscle surface mesh and moment arm. The task is formulated as a least-squares optimization problem with a threefold objective for the path to: (1) pass through multiple ellipses representing muscle cross-sections, (2) yield moment arms that match experimental measurements, and (3) yield moment arms in the expected functional directions. We demonstrate the performance of our optimization framework with the musculoskeletal surface mesh from the Visible Human Male and moment arm datasets from literature—producing 42 paths that are anatomically realistic and biomechanically accurate within 20.1 minutes. Our optimization framework is gradient-specified, which is faster and more accurate than using the default numerical gradient, making it applicable for large-scale subject-specific uses.

## 1. Introduction

In musculoskeletal modeling, a musculotendon path is a simplification of the 3-D musculotendon geometry, and often it is a cable that reflects the *shape* of a musculotendon unit^1^ as well as its tendinous *attachments* on the bones (Garner and Pandy, 2000; Hicks et al., 2015; Modenese and Kohout, 2020). Depending on how simplified the musculotendon geometry needs to be, the path is configured based on anchor points (i.e., the origin, via, and insertion points) and wrapping obstacles (e.g., cylinders): The simplest path is a straight cable defined by the origin and insertion, while via points and wrapping obstacles may be used for extra geometric features such as turnings and curvatures (Delp et al., 1990; Garner and Pandy, 2001; Ma’touq et al., 2019). Importantly, the locations of the anchor points (and obstacles) are not defined with respect to the same bone, so the path geometry will change due to joint rotation as the musculotendon geometry does.

Despite such geometric simplification, a well-modeled path should retain two properties of the muscle across different joint configurations, namely its length and its moment arm (Garner and Pandy, 2001; Hicks et al., 2015). These are crucial properties that the accuracy of musculoskeletal simulations depends on: In a forward simulation, given the neuromuscular signal, muscle length is related to how much force can be exerted (Guo et al., 2022; Millard et al., 2024), and muscle moment arm determines how much joint moment (hence what motion and how much end-effector force) is to be generated from this force (McFarland et al., 2023; Ong et al., 2016; Zajac, 1993). In an inverse simulation, given the dynamic motion, muscle lengths and moment arms (especially the latter) are the key to estimating joint reaction forces and muscle activations (Beaucage-Gauvreau et al., 2019; Erdemir et al., 2007; Hamner and Delp, 2013; Karabulut et al., 2020; Prilutsky and Zatsiorsky, 2002). Therefore, the mission of muscle path modeling is in some sense to replicate the length–joint angle relation as well as the moment arm–joint angle relation of the target muscle.

In general, there are two approaches to replicate these relations. One is the *cause*-oriented approach, where the bones and adjacent tissues are regarded as the cause of muscle deformation (Lloyd et al., 2021; Wang et al., 2025). In other words, the wrapping obstacles and via points should represent specific anatomical structures. The assumption is, with the right origin and insertion, so long as the anatomical obstructions are correctly reflected, the paths should deform at appropriate joint configurations as real muscles do; accordingly, the length– and moment arm–joint angle relations should be accurate. On the other hand, if length or moment arm data are available, they can be directly used to calibrate the muscle path as an *effect*-oriented approach (Chen et al., 2026; Livet et al., 2022; Modenese and Kohout, 2020). That is, the length– and moment arm–joint angle relations are regarded as the effect of muscle deformation. Regardless of whether the anatomical obstructions are reflected in the muscle path, as long as the key relations are replicated, the simulations should be accurate.

In Chen et al. (2026), we demonstrated an automated method of effect-oriented muscle path calibration: With artificial moment arm data from a reference model as the target value, a muscle path is configured to yield almost the same moment arm–joint angle relation as in the reference path. The process of parameter tuning for matching moment arms is formulated as a least-squares optimization problem, and our method features specifying the gradient of the cost function in its analytical form, which enables efficient computation and rapid convergence compared with using the numerical gradient.

Nevertheless, there are two issues remaining to be accounted for. First, although we succeeded in replicating the high-dimensional moment arm–joint angle relation using *in silico* data (Chen et al., 2026), it is crucial that the method remains effective with experimental measurements, which are generally less comprehensive. The moment arm–joint angle relation is high-dimensional in the sense that a muscle may actuate multiple degrees of freedom (DoF): It may have multiple moment arms (McCullough et al., 2011; Vaarbakken et al., 2014; Wretenberg et al., 1996), and each moment arm is susceptible to the motion in each of the relevant DoFs (Delp et al., 1999; Hintermann et al., 1994; Wolfram et al., 2018). However, moment arm measurement is generally limited to the dominant DoF (e.g., ankle plantarflexion for the triceps surae or knee flexion for the hamstring), and the available moment arm–joint angle relation is mostly a curve (Chen and Franklin, 2025). Moreover, small muscles such as the gluteus minimus, the pectineus, and the gemelli have no moment arm measurement at all (Chen and Franklin, 2025). It is not entirely clear how our method would perform based only on the dominant moment arms, and alternative sources of data are needed to model muscles lacking moment arm measurements. Second, if moment arm is the only property to be calibrated, then length may not be well matched, and the resultant muscle path may be partially unrealistic in appearance. Notice that muscle moment arm is the partial derivative of muscle length with respect to the correspondent joint angle (An et al., 1984; Sherman et al., 2013), and matching muscle moment arm only guarantees matching the derivative of muscle length, not the property itself.

With regard to these issues, it is important that morphological information is brought into the optimization problem; e.g., MR images of the muscles. Such information is useful in two perspectives: First, with 3-D mesh reconstructed from medical images, we may confine the muscle path to pass through the mesh volume, making it appear anatomically realistic and have its length fall in a reasonable range. Second, for muscles with few or no moment arm measurements, the geometric features as well as the alignment relative to the bones are a valuable indication of how the moment arms should be (An et al., 1984; Herzog and Read, 1993). Recently, Modenese and Kohout (2020) proposed a method of automated path modeling based on musculotendon surface mesh: By extracting cross-sections from the 3-D mesh, one or multiple paths can be defined by connecting anchor points defined at the same relative positions in each cross-section (imagine the hollow pathways in a lotus root). Essentially, this method aims to match muscle length, and the paths modeled in this fashion will appear anatomically realistic; yet the moment arm–joint angle relations are not necessarily correct. Although they manipulate the kinematics of the anchor points to avoid aberrant path deformation, the related parameters are not tuned based on muscle length or moment arm. In other words, the visual resemblance of the path does not guarantee biomechanical accuracy across a vast range of motion (RoM).

In this work, we develop a hybrid calibration based on muscle surface mesh and moment arm for path modeling in the lower limb. Rather than focusing on a single source of data, we utilize surface mesh as a confinement and place additional emphasis on moment arm calibration. The goal is to have the path retain sufficient geometric features while yielding moment arms that match experimental measurements and remain consistent with known muscle functions. An overview is shown in Fig 1 for the workflow of the proposed method.

**Figure 1.**
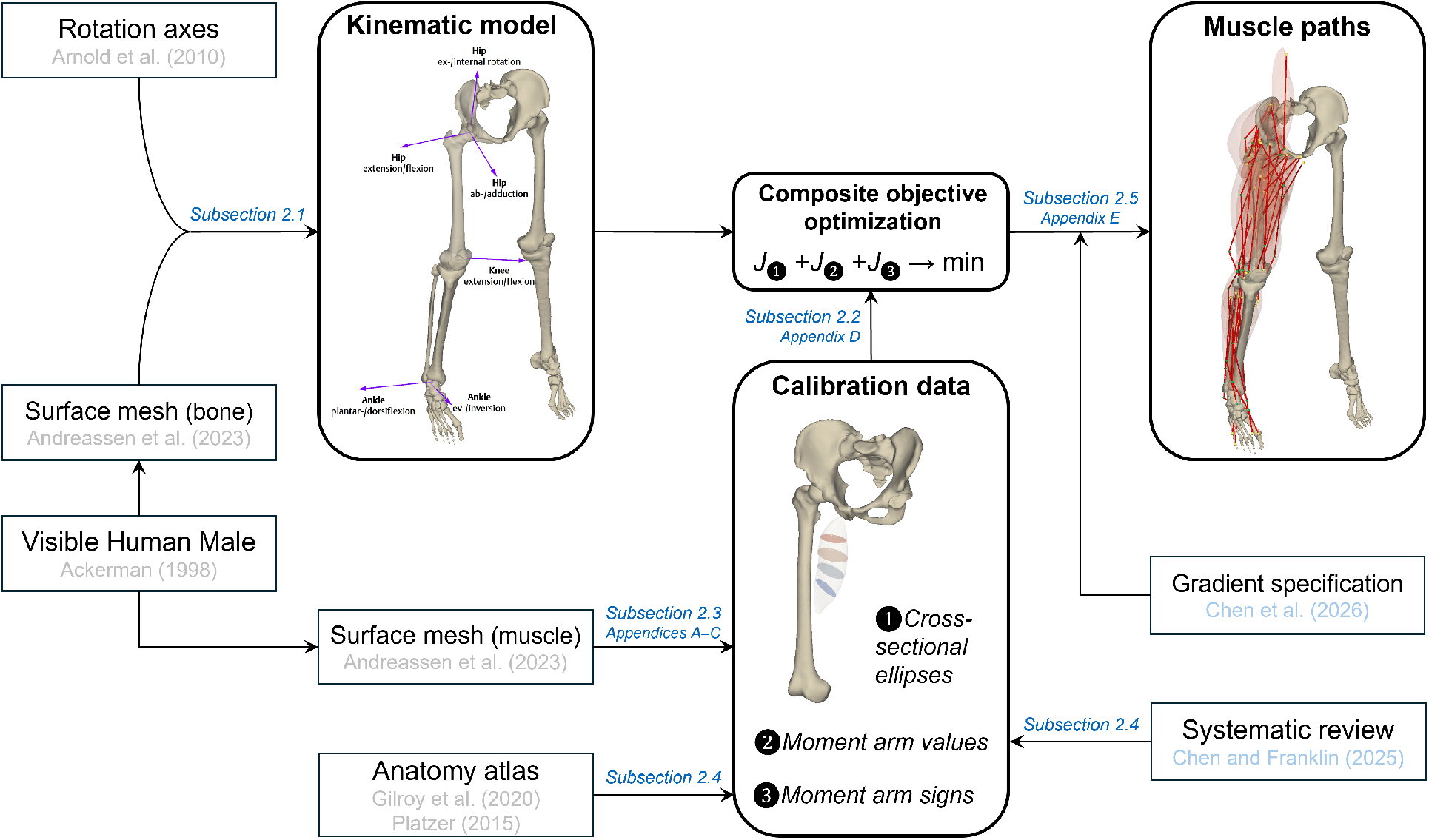
Workflow of the proposed method.

## 2. Methods

### 2.1. Kinematic Model

As a prerequisite for muscle path modeling, a 6-DoF lower-limb kinematic model is developed with the bone surface mesh from the Visible Human Male (VHM) dataset (Ackerman, 1998; Andreassen et al., 2023) and the kinematics replicated from the Arnold lower-limb model (Arnold et al., 2010). The six DoFs include:

1. hip extension/flexion,
2. hip ab-/adduction,
3. hip ex-/internal rotation,
4. knee extension/flexion,
5. ankle dorsi-/plantarflexion, and
6. ankle ever-/inversion.

The sign rule for angle and moment arm in each DoF is based on the ISB recommendations (Derrick et al., 2020; Wu et al., 2002) except for knee extension/flexion, which is reversed so that moment arms for anti-gravity motions share the negative sign.

For each rotation, the center is manually determined on the VHM skeleton to visually match the relative position in the Arnold model, while the axis has the same direction as defined in the Arnold model; exceptions are discussed in the *Joint Kinematics* subsection. Since the VHM skeleton is obtained from a cadaveric specimen (Ackerman, 1998), it is posed in the fixation position, and we manually adjusted each DoF to pose it into a neutral position visually similar to the default position of the Arnold model. The joint configuration for the VHM fixation position is defined as 1.5^°^ hip extension, 6^°^ hip abduction, 10^°^ hip internal rotation, 0^°^ knee flexion, 25^°^ ankle plantarflexion, and 30^°^ ankle inversion. Importantly, we also follow the convention in both OpenSim (Delp et al., 2007) and the Arnold model for the Euler angle sequences. This way, given any joint configuration, our model will be posed the same as the Arnold model does in OpenSim.

### 2.2. Composite objective optimization

Muscle path modeling is formulated as a least-squares problem with the composite objective cost function

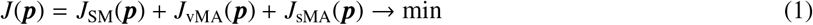

where ***p*** denotes path-related parameters, and *J*_SM_(***p***), *J*_vMA_(***p***), and *J*_sMA_(***p***) are the cost terms related to surface mesh and moment arm. The indications of these terms will be further explained in the following subsections. As a brief summary, the threefold objective is to have the muscle path

1. pass through multiple ellipses that represent cross-sections of the muscle,
2. yield moment arms that match experimental measurements, and
3. yield moment arms with the correct signs (i.e., functional directions).

We use the obstacle-set method (Garner and Pandy, 2000) to configure the muscle path. The composition of ***p*** and other details regarding the path configuration are explained in detail in Chen et al. (2026). Here, a muscle may be simplified as a straight path (OI), a path with one via point (OVI), a path with two via points (OV_1_V_2_I), a path with one cylinder as wrapping obstacle (OCI), or a path with two cylindrical obstacles (OC_1_C_2_I). In terms of the points, a muscle path may come in one of the following form:

1. O − I,
2. O − V − I,
3. O − V_1_ − V_2_ − I,
4. O − W_C_ ⌢ W_C_′ − I,
5. O − W_C_1 ⌢ W_C_′1 − W_C_2 ⌢ W_C_′2 − I;

where O and I are the origin and insertion points, and V and W denote the via and wrapping points; if wrapped, each cylindrical obstacle (C) produces at most one pair of wrapping points. Accordingly, a path can be divided into one, two, three, or six sub-segments, indicated by “−” (straight) and “⌢” (curved).

The reference frames for the anchor points and obstacles are predetermined based on the anatomy (Gilroy et al., 2020; Platzer, 2015). For instance, the origin and insertion points of the gastrocnemius lateralis are fixed respectively on the femur and the calcaneus. If a single via point (or cylinder) is required to reach the objectives, it will be fixed on the tibia. If a second via point (or cylinder) is required, its reference frame depends on the muscle; in the case of the gastrocnemius, it will be fixed on the tibia, since the muscle spans mostly on this bone.

As we recently demonstrated in Chen et al. (2026), gradient-specified optimization is advantageous in terms of both speed and accuracy compared with when the gradient is computed numerically. For the analytical gradient to exist, the cost terms in Eq. 1 must be differentiable across the entire domain. In the following subsections, we will show how to structure these terms in a continuous form by replacing the conditional statements and nondifferentiable components with soft functions.

### 2.3. Surface Mesh

With the morphological information from the surface mesh, we aim to limit the muscle path from forming aberrant routes, and the path length will roughly match the muscle length. In this study, we use the muscle meshes from Andreassen et al. (2023) for both their easy accessibility and their compatibility with our kinematic model. As is with the bone meshes, the muscle meshes are also obtained in the fixation position.

There are multiple ways to confine a path to the surface mesh. For example, if the tendon mesh is included along with the muscle mesh, we can determine some centerline for the entire mesh, and directly set it as the path. However, this leaves little room for the path to be tuned for moment arm calibration, and for many muscles, the tendon is absent in Andreassen et al. (2023)—so the mesh centerline no longer represents the entire muscle-tendon unit. Therefore, we compute multiple ellipses that represent cross-sections of the muscle belly, and the objective is set so that the path pass through each ellipse. This way, the path will still form the necessary turnings and curvatures as the muscle does, and it can be adjusted of the moment arms while staying within the mesh volume.

The relative position between the path and the mesh is determined by computing path intersection with each of the *N*_ell_ cross-sectional ellipses. The cost term for surface mesh is structured based on the number of unintersected ellipses

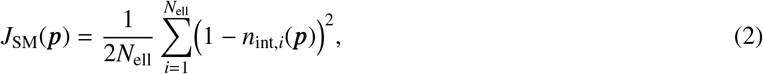

where *n*_int_ denotes the boolean for valid intersection, and the term 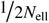is a weight factor for cost normalization (see Appendix D). Notice there are ways to determine the positional relativity directly with the mesh vertices, but the use of ellipses reduces the computational cost significantly with little impact on the results.

Figure 2 shows how the ellipses are obtained from a muscle surface mesh. A key is to identify two vertices on the surface mesh directing the entry and exit for the path (Fig 2, left). This can be done manually for each muscle, but in the spirit of automation, here we use the pseudocode in Appendix A to find such vertices. With them as boundary constraints, a harmonic scalar field is constructed over the muscle mesh (Dong et al., 2005): Each vertex is assigned a value between 0 and 1 depending on its relative position between the entry and exit points (Fig 2, mid-left). Based on the field isolines, we can then extract a series of vertex sets (Fig 2, middle), each of which will be fitted with an ellipse as an approximate estimation of a cross-section (Fig 2, mid-right; see Appendix B).

**Figure 2.**
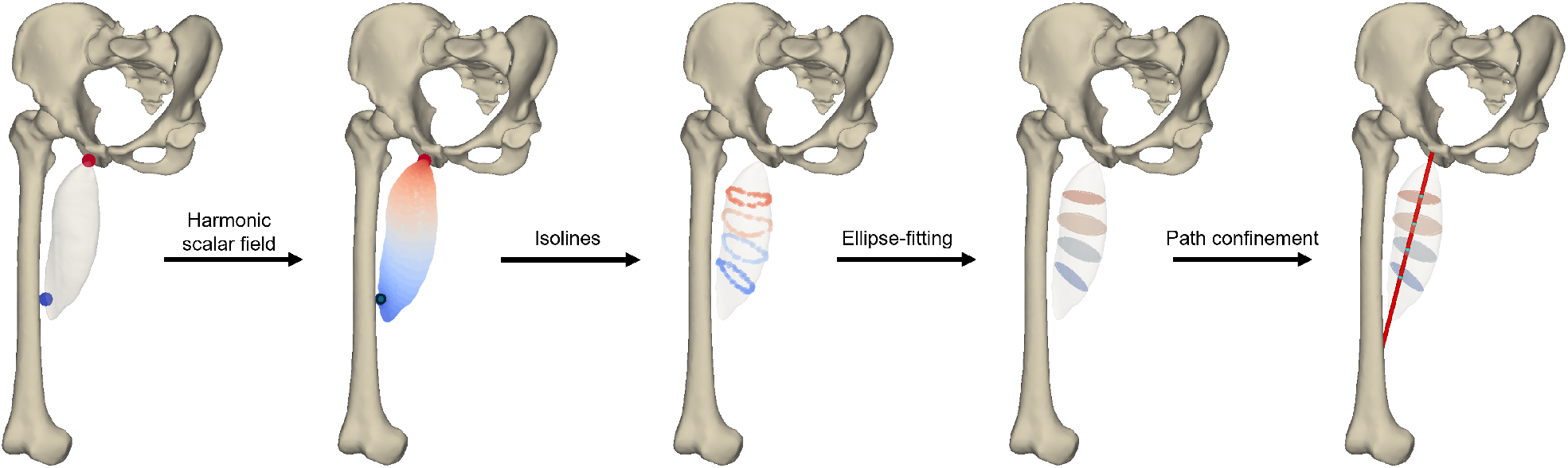
Muscle surface mesh as a confinement for the path (exampled by the adductor longus). Left: Identify two vertices on the muscle mesh directing the entry (red) and exit (blue) of the path. Center-left: Construct a harmonic scalar field over the mesh with the entry and exit points as boundary constraints—a scalar value between 0 and 1 (indicated by the color) is assigned to each vertex. Center: Extract a number of vertex sets based on the field isolines. Center-right: Fit an ellipse to each vertex set in representation of a muscle cross-section. Right: The path is expected to intersect all cross-sectional ellipses.

A path is considered to pass through the muscle mesh if each cross-sectional ellipse is intersected by at least one of the sub-segments (Fig 2, right). As described in Appendix C, for each sub-segment, we first compute its intersection point on the plane in which the ellipse lies and the scalar indicator *s*_1_ describing the relative location of the intersection point on the sub-segment. Then we compute a second scalar indicator *s*_2_, which describes the location of the intersection point relative to the ellipse.

Importantly, the intersection is valid if two conditions are satisfied: 0 < *s*_1_ < 1, and *s*_2_ > 0. The first condition ensures it is the sub-segment itself (rather than its extended line) that intersects the ellipse, and the second ensures it is the ellipse itself (rather than its expanded space) that is intersected by the sub-segment. In combination, this twofold conditional statement contains four branches, only one of which leads to a valid intersection. For the optimization, we expect it to find the “valid” branch without getting trapped in the “invalid” branches.

Take the example of when 0 < *s*_1_ < 1, we have *n*_int_ = 1 if *s*_2_ > 0, and *n*_int_ = 0 if *s*_2_ < 0. In both cases, ∂*n*_int_/∂***p*** = 0, and *n*_int_(***p***) is not differentiable at *s*_2_ = 0. This means that, apart from returning NaN at *s*_2_ = 0, such a gradient might not direct the optimization to proceed from the domain where *s*_2_ < 0 to where *s*_2_ > 0—reflected as a path sub-segment being adjusted from outside an ellipse to inside. The same issue exists as *s*_1_ tends to 0 and 1.

To address this issue, we use a soft ReLU function to modify the quadruple-branched *n*_int_ into a smooth form

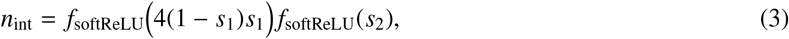

where

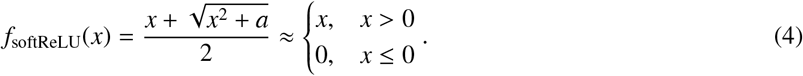

Here, *a* should be a sufficiently small number (e.g., 0.01). If *s*_1_ < 0, *s*_1_ > 1, or *s*_2_ < 0, the input for *f*_softReLU_ is negative, and *n*_int_ tends to 0; otherwise, *n*_int_ is positive and tends to 4(1−*s*_1_)*s*_1_ *s*_2_, whose maximal value is approximately 1.

The validity of intersection is evaluated pairwise between all ellipses and all sub-segments (*N*_ell_×*N*_seg_). A subsegment may intersect multiple ellipses, but if an ellipse is intersected by multiple sub-segments, only one is counted. For this, we have

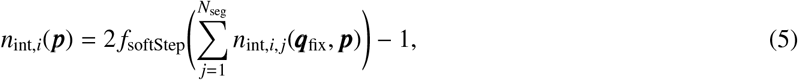

where ***q***_fix_ is joint configuration for the VHM fixation position and the subscripts indicate the evaluation is between the *i*-th ellipse and *j*-th sub-segment, and

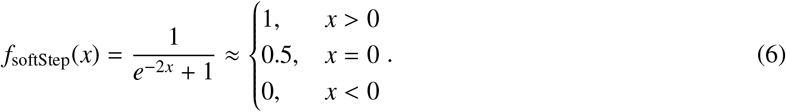

If an ellipse is intersected (regardless how many times), the input for *f*_softStep_ is positive, and its output approaches 1. In the case of no intersection (or if the input is sufficiently small), the soft function yields approximately 0.5 instead of 0, thus it is normalized in Eq. 5 so that *n*_int_ ∈ (0, 1) .

In Eq. 2, *J*_SM_ tends to 0 if each ellipse is intersected by at least one sub-segment. Since *n*_int_ is smoothed with *f*_softStep_ and *f*_softReLU_, the analytical form of ∂*n*_int_/∂***p*** can be derived using the chain rule, then we may specify the gradient for *J*_SM_(***p***). See an example of the mathematical derivation of gradients in Chen et al. (2026).

### 2.4. Moment Arm

If measurements are available for any moment arm of a muscle, then we simply refer to our recent work for the calibration (Chen et al., 2026). For example, we can evaluate if the model moment arm falls in a certain range of the measurement with

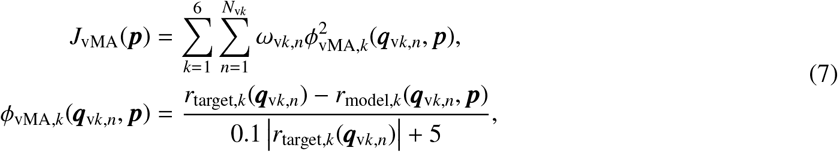

where *r*_target,*k*_ and *r*_model,*k*_ denote the target value and the model output for the *k*-th moment arm (with the unit of mm) at joint configuration ***q***_v*k,n*_, and ω_v*k,n*_ is a weight factor (see Appendix D). Apparently, *ϕ*_vMA,*k*_ depends on the ratio of the difference between *r*_target,*k*_ and *r*_model,*k*_ with respect to the tolerance of (10% |*r*_target,*k*_| + 5) mm.

In preparation for the present study, we have previously summarized 300 moment arm datasets in the hip, knee, and ankle (Chen and Franklin, 2025). However, for many muscles, there are only a few measurements or even no measurements at all. In this case, the most we can do is to designate a moment arm to carry the sign indicated by the anatomy. For example, the adductors longus, brevis and magnus are obviously hip adductors, and as an easy objective, the paths should yield a positive moment arm in the sagittal plane of the hip. This can be achieved with

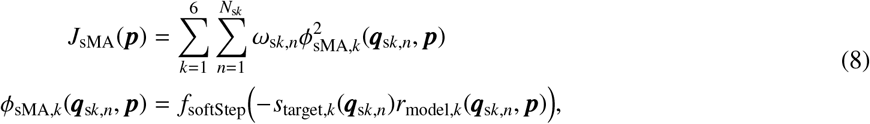

where *s*_target,*k*_ is the target sign for the *k*-th moment arm at joint configuration ***q***_s*k,n*_, and ω_s*k,n*_ is a weight factor (see Appendix D). When the sign of *r*_model,*k*_ matches *s*_target,*k*_, the input for *f*_softStep_ is negative, and ϕ_sMA,*k*_ tends to 0; otherwise, it tends to 1.

As a focus of our recent work, we derived the analytical form of ∂***r***_model_/∂***q*** (Chen et al., 2026). With the chain rule, we may easily specify the gradients for both *J*_vMA_(***p***) and *J*_sMA_(***p***) without much extra work.

### 2.5. Calibration

With the analytical form of ∂*J*/∂***p***, we used the nonlinear least-squares solver (lsqnonlin) in MATLAB to solve Eq. 1. Notice the outcome of Eq. 1 is affected by the composition of ***p***, which depends on how the path is structured. To find the best possible solution, the calibration was programmed to iterate through each path structure unless the optimization cost reaches a sufficiently small level (e.g., 0.03): For any muscle, we start with OI—the most simple structure—and incrementally increase structural complexity if necessary, progressing in the order of OVI, OV_1_V_2_I, OCI, and OC_1_C_2_I. Then, from all available solutions, the final solution is selected considering the trade-off between calibration accuracy and structural simplicity. More complex path structures incur higher cost penalties: 1.25 for OVI, 1.5 for OV_1_V_2_I, 2 for OCI, and 4 for OC_1_C_2_I.

The optimization was performed on a 2.9-GHz Intel Core i9 with 64 GB RAM and 14 CPU cores using parallel computing (`parfor`). For each path structure, we arranged parallel computing to optimize 14 initial points simultaneously, with the aim to reduce the risk of local minimum. See Appendix E for other configuration details and the generation of the initial points.

For surface mesh calibration, we extracted four ellipses from each muscle mesh. For moment arm calibration, we selected appropriate measurements from the literature as the target values (*r*_target_) and determined muscle functions based on Gilroy et al. (2020) and Platzer (2015) for the target signs (*s*_target_). See Appendix F for the target values and target signs in detail. All calibration data can be found in Supplementary Materials.

## 3. Results

The calibration of 42 muscle paths was completed in 20.1 min, and the configurations of the resultant paths are visualized in Figs. 3–5. For 29 paths, the iteration over potential structures terminated in advance by reaching below the cost threshold, and the final structures were either straight or via point–based. For the other 13 paths, although all five structures were iterated, none of the final structures were obstacle-based. Overall, 10 paths were structured as OI, 20 paths as OVI, and 12 as OV_1_V_2_I.

**Figure 3.**
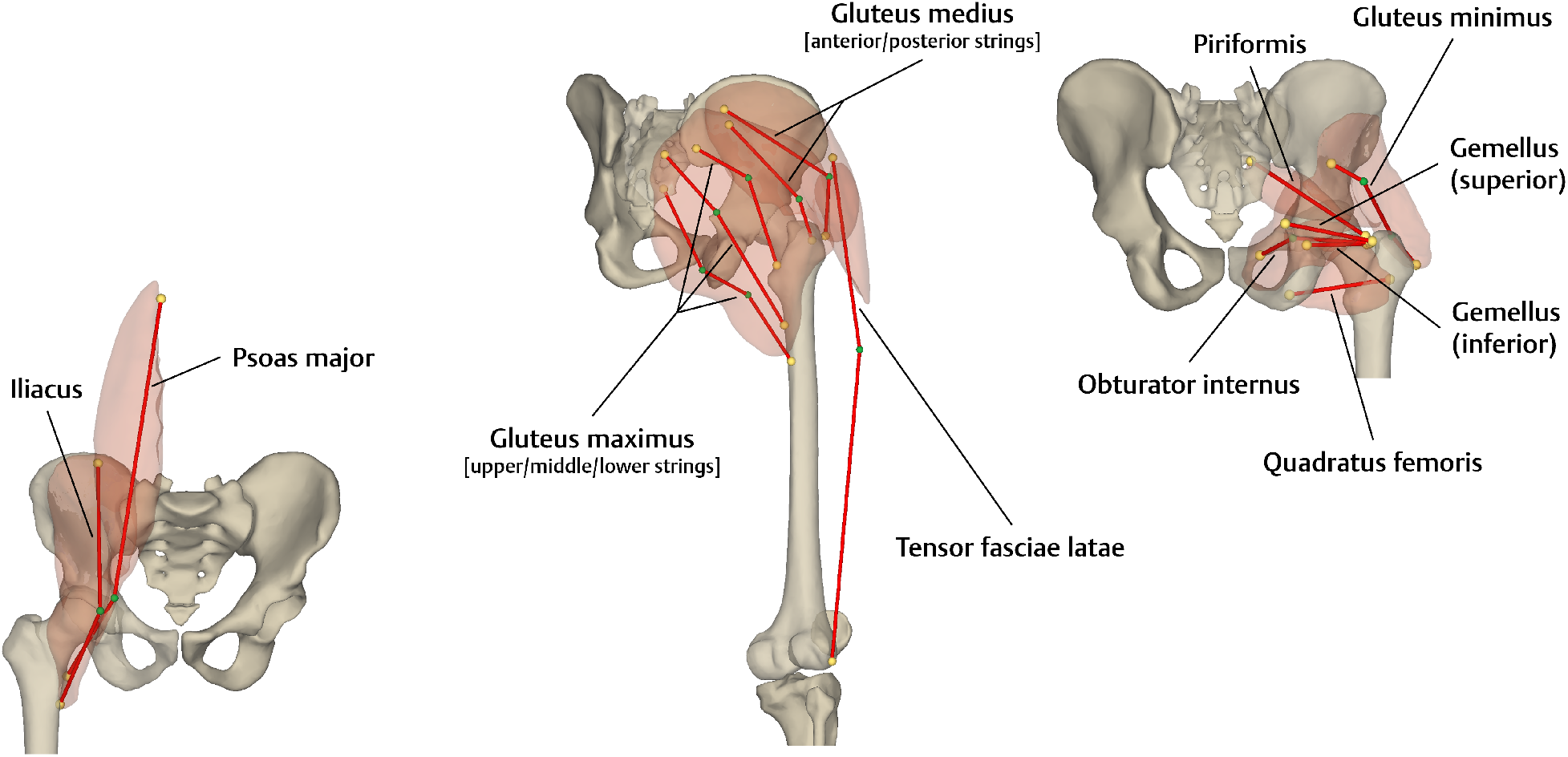
Path configurations for the iliopsoas and the gluteal muscles. Left: Iliopsoas. Center: Superficial gluteal muscles. Right: Deep gluteal muscles. The origin and insertion points are colored in yellow, and the via points are in green.

**Figure 4.**
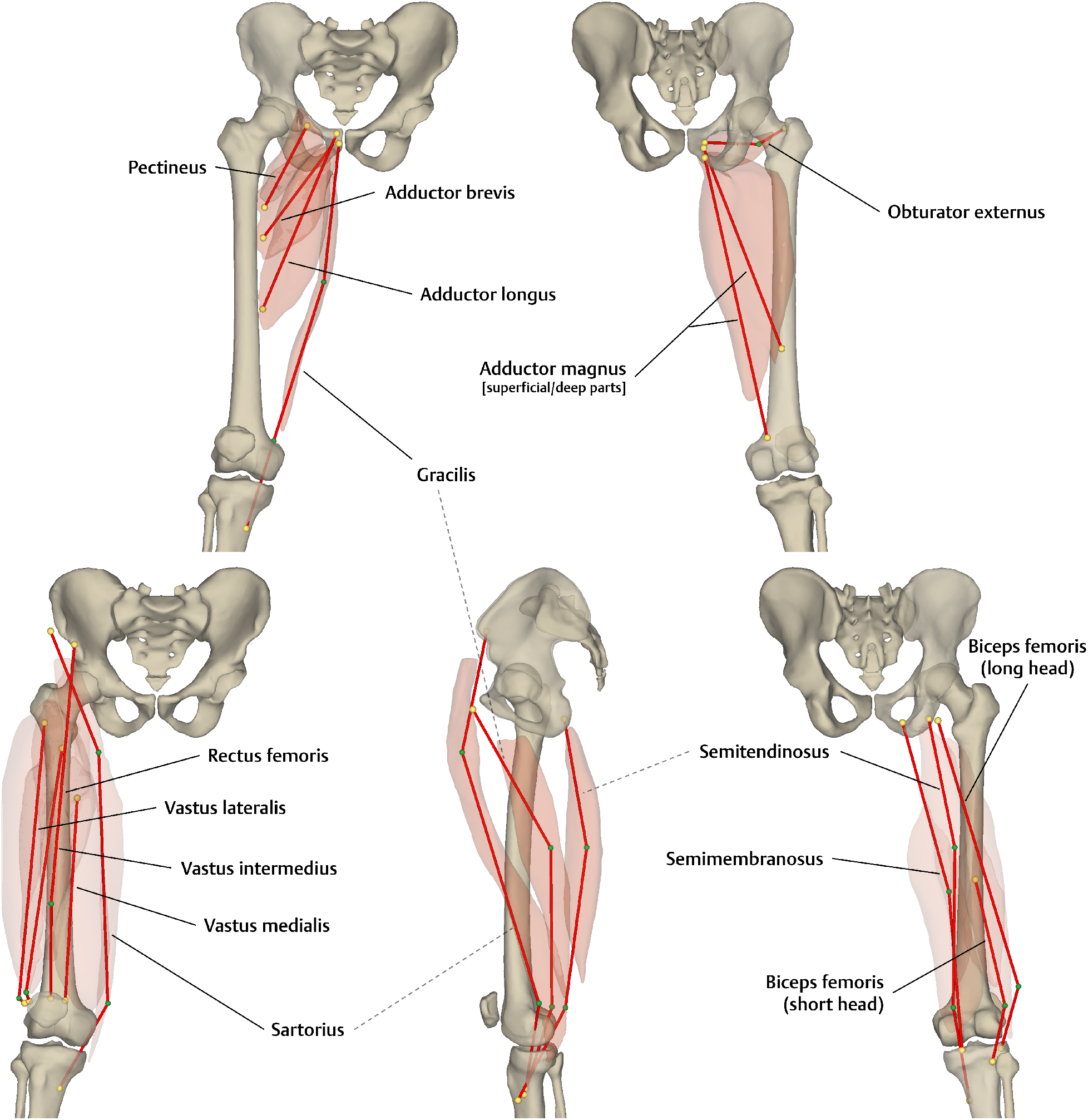
Path configurations for the thigh muscles. Top left: Superficial layer of medial compartment. Top right: Deep layer of medial compartment. Bottom left: Anterior compartment. Bottom center: Muscles that co-insert into the pes anserinus. Bottom right: Posterior compartment (i.e., hamstrings). The origin and insertion points are colored in yellow, and the via points are in green.

**Figure 5.**
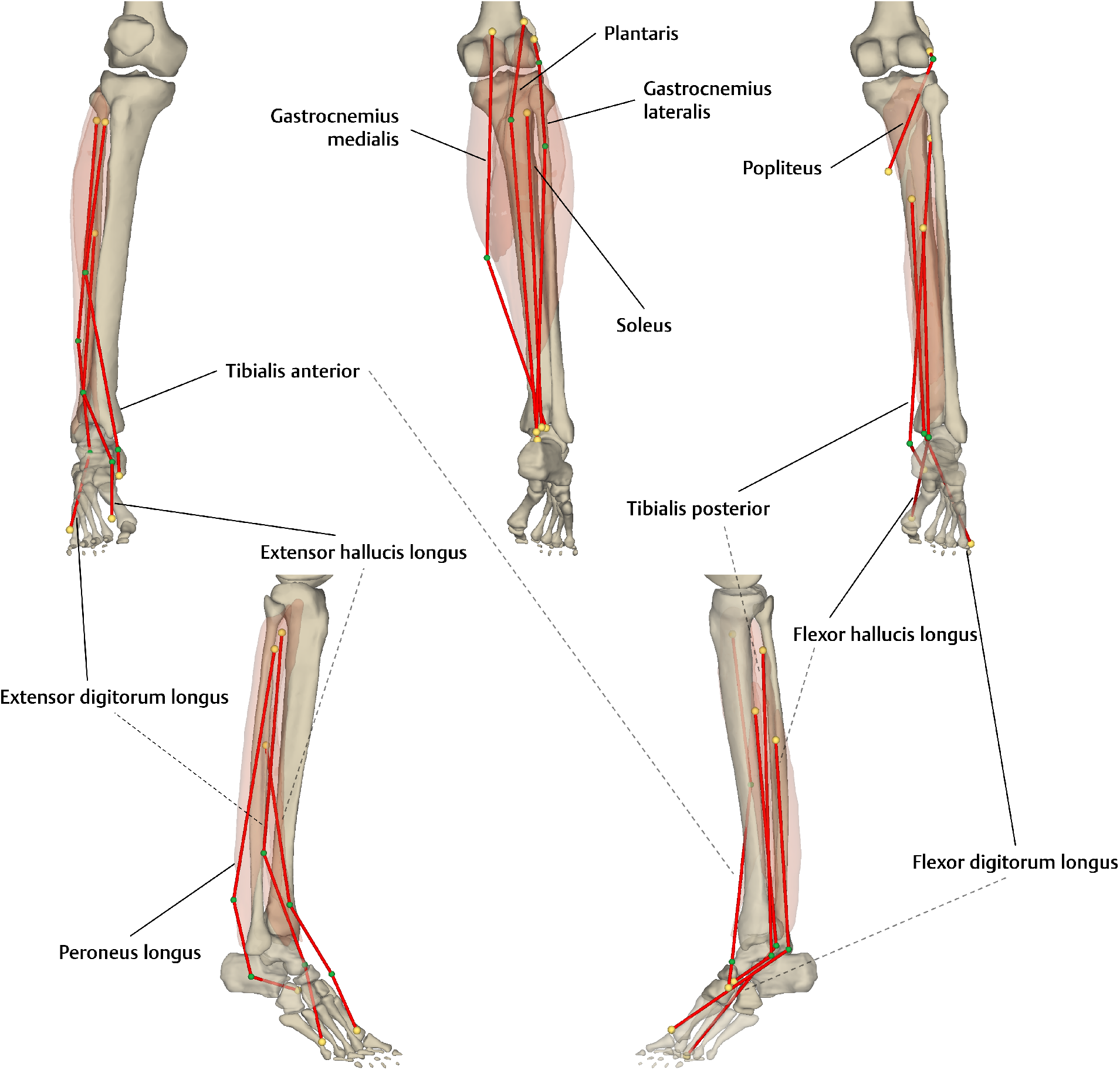
Path configurations for the leg muscles. Top left: Anterior compartment. Top center: Superficial layer of posterior compartment. Top right: Deep layer of posterior compartment. Bottom left: Muscles that insert into the lateral foot. Bottom right: Muscles that insert into the medial foot. The origin and insertion points are colored in yellow, and the via points are in green.

The cost and time of calibration as well as the final structure for individual paths are reported in Appendix F. Other details regarding the calibration process and results can be found in Supplementary Materials. Representative cases are discussed in the next section.

## 4. Discussion

### 4.1. Muscle with Sufficient Moment Arm Data

As we reported in Chen and Franklin (2025), available moment arm measurements are mainly found in the sagittal plane, and it is rather difficult to gather sufficient data for the moment arm–joint angle relations across all actuating DoFs of a muscle. For instance, to accurately calibrate a monoarticular muscle across the 3-DoF hip joint, we need measurements of three moment arm relations, each of which have three dependent variables (i.e., the joint angle in each of the three DoFs); this is equivalent to three 3-D heatmaps and so far no such data exist.

In our kinematic model, the knee is simplified as a 1-DoF joint as in Arnold et al. (2010), so there is a lower demand for the calibration data of its monoarticular muscles, such as the vasti, the short head of the biceps femoris, and the popliteus. Figure 6 shows the results for these muscles: With the moment arm–joint angle relation being a single curve, the path is easily aligned within the muscle mesh, and there is no risk of potential inaccuracy in the hip and ankle DoFs.

**Figure 6.**
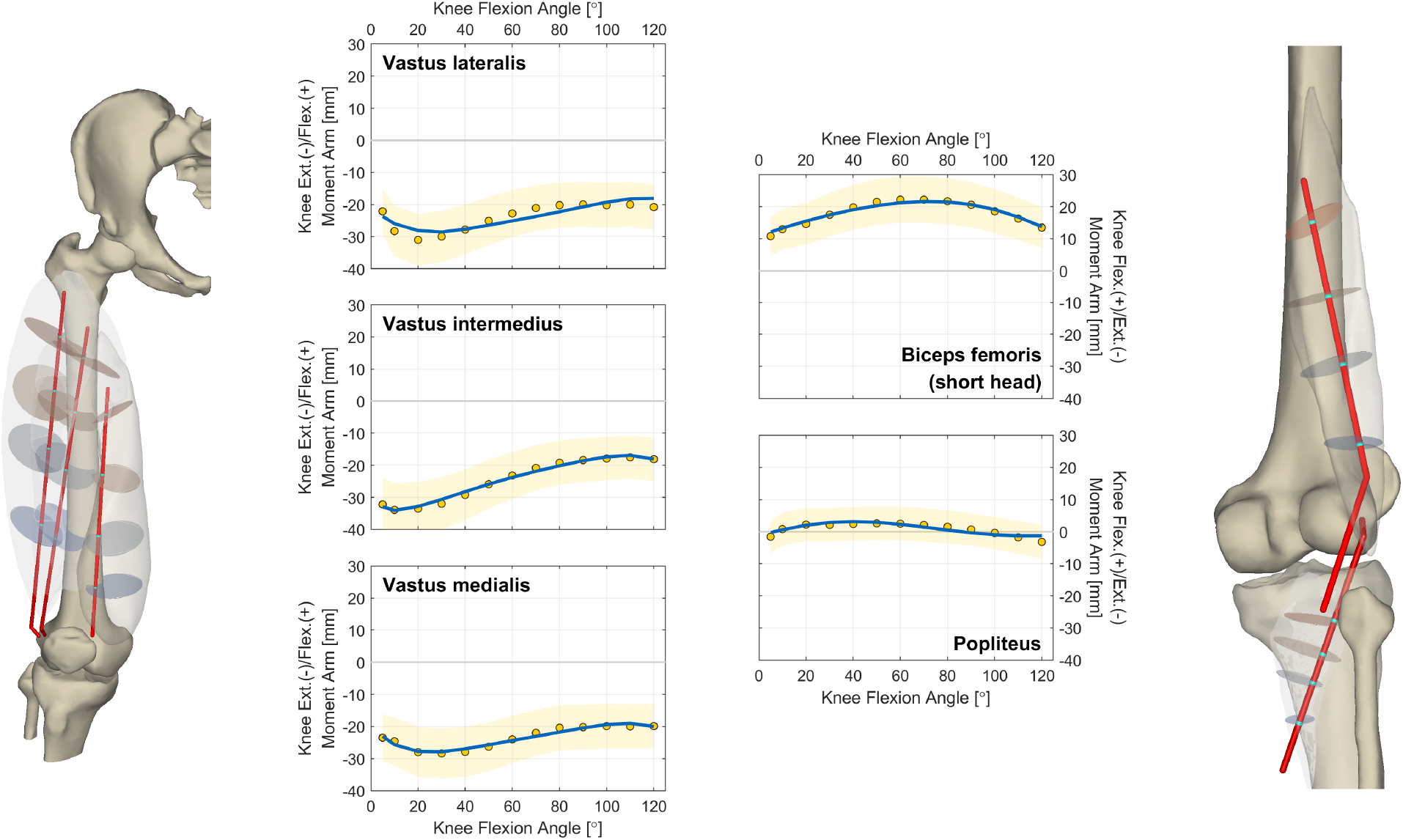
Calibration results of monoarticular muscles in the knee. Left and Center-left: Ellipse–path intersections and 1-D moment arm relations of the extensors. Center-right and Right: 1-D moment arm relations and ellipse–path intersections of the flexors. In the models, each cyan node along the path denotes a valid intersection with one of the ellipses. In the charts, the blue curves depict the model moment arms, the yellow scatters depict measurements from Buford et al. (1997), and the yellow areas represent the tolerance ranges for moment arm calibration.

### 4.2. Muscle with Limited Moment Arm Data

For muscles with more actuating DoFs, there are more practical issues. Take the example of the soleus, a major ankle plantarflexor with a role in ever-/inversion. In theory, its plantarflexion moment arm is dependent on both the dorsi-/plantarflexion and ever-/inversion angles; in other words, the relation is depicted by a surface. In our case, due to data scarcity, the calibration is based on only two curves, one with the dorsi-/plantarflexion angle as the variable (while ever-/inversion is neutral) and another with varying ever-/inversion (neutral dorsi-/plantarflexion); see Fig. 7 (top right, blue curves). Although these two “orthogonal” curves form the “backbone” of the 2-D relation, it is still possible that the plantarflexion moment arm is inaccurate in other joint configurations; e.g., when the ankle is dorsiflexed while inverted. To effectively calibrate such a moment arm relation, the ideal data would be measurements spreading across the ankle RoM, or should at least cover joint configurations involved in the musculoskeletal simulation. A similar issue exists with the ever-/inversion moment arm of the soleus, where only three curves are available for the calibration of a surface (Fig. 7, bottom right).

**Figure 7.**
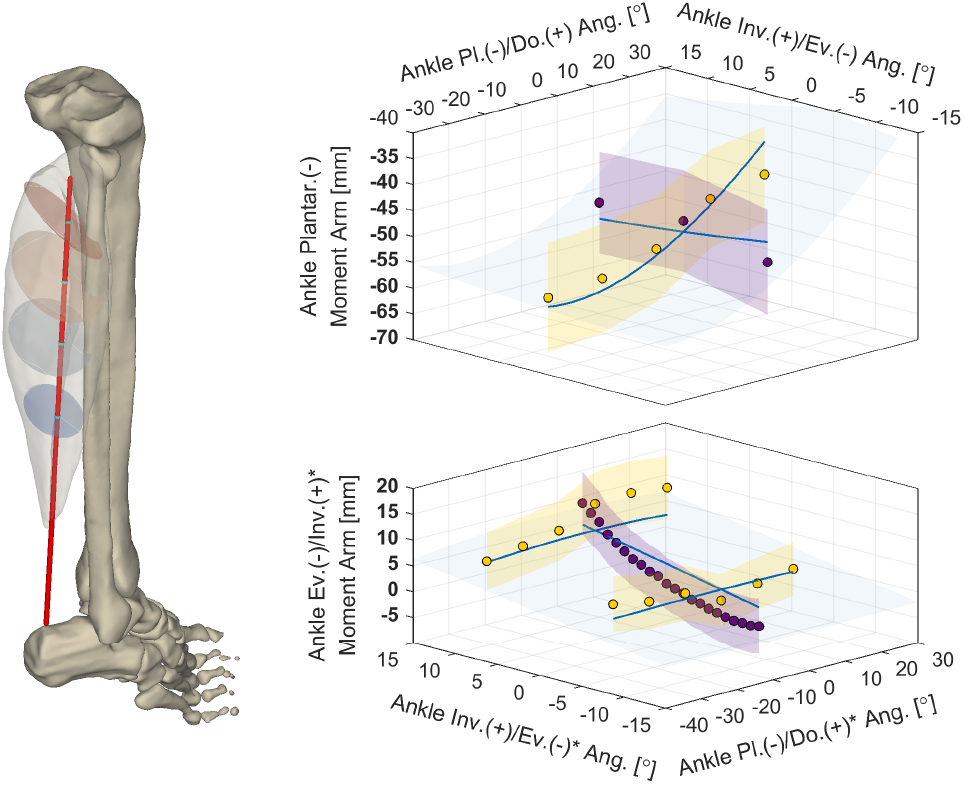
Calibration results of the soleus. Left: Ellipse–path intersections (denoted by the cyan nodes). Top right: 2-D relation of ankle plantarflexion moment arm with ankle angles; in comparison with measurements from Maganaris et al. (2000) and Wolfram et al. (2018), depicted by the yellow and purple scatters. Bottom right: 2-D relation of ankle eversion/inversion moment arm with ankle angles; in comparison with measurements from McCullough et al. (2011) and Hintermann et al. (1994), depicted by the yellow and purple scatters. The asterisks in the axis labels denote that the kinematics are slightly different (see the *Joint Kinematics* subsection). In the 3-D charts, the blue surfaces depict the model moment arms, the yellow and purple areas represent the tolerance ranges for moment arm calibration, and the blue curves denote the intersections between the blue surface and the yellow/purple areas, depicting the 1-D moment arm relations in the correspondent joint configurations.

A key message here is that, as the moment arm–joint angle relation gets more complex, the amount of data required for calibration increases drastically. If the available data are limited, it is difficult to tell if the uncalibrated part of the moment arm relation is accurate. In particular, when an uncalibrated configuration is far from the calibrated ones in the joint space, there is a higher risk of inaccuracy—in the worst case, the muscle may even function as an antagonist. For muscles lacking moment arm measurements, we have the sign-based cost term (Eq. 8) to prevent them from becoming antagonists in the designated joint space. Take the example of the lower part of the gluteus maximus, which is a hip extensor, adductor, and external rotator. As shown by the 3-D heatmaps in Fig. 8 (top right and bottom), each of its three moment arms are theoretically dependent on the angles in all three DoFs of the hip. However, for each moment arm, the available dataset is only 1-dimensional, containing measurements with one angle as the variable, shown as the colored scatters in Fig. 8. Calibration based on these data has no demand on the muscle functions as in other joint configurations, so we need to establish additional rules. For instance, considering the relative position between the gluteus maximus and the femur head, it is hard to imagine this muscle functioning as a hip flexor at all, so we designated the hip extension/flexion moment arm sign to remain negative in any joint configuration. In Fig. 8 (top right), the slices of 3-D heatmaps are mostly blue (indicating a negative moment arm), and the little joint space where the moment arm is great than −5 mm (the path being too close to the rotation center) is marked out in gray. For the other two moment arms, it is unclear within what RoM the lower part of the gluteus maximus contributes to hip adduction and external rotation. Therefore, we did not designate the moment arm signs for a series of extreme joint configurations, and the results are shown in Fig. 8 (bottom).

**Figure 8.**
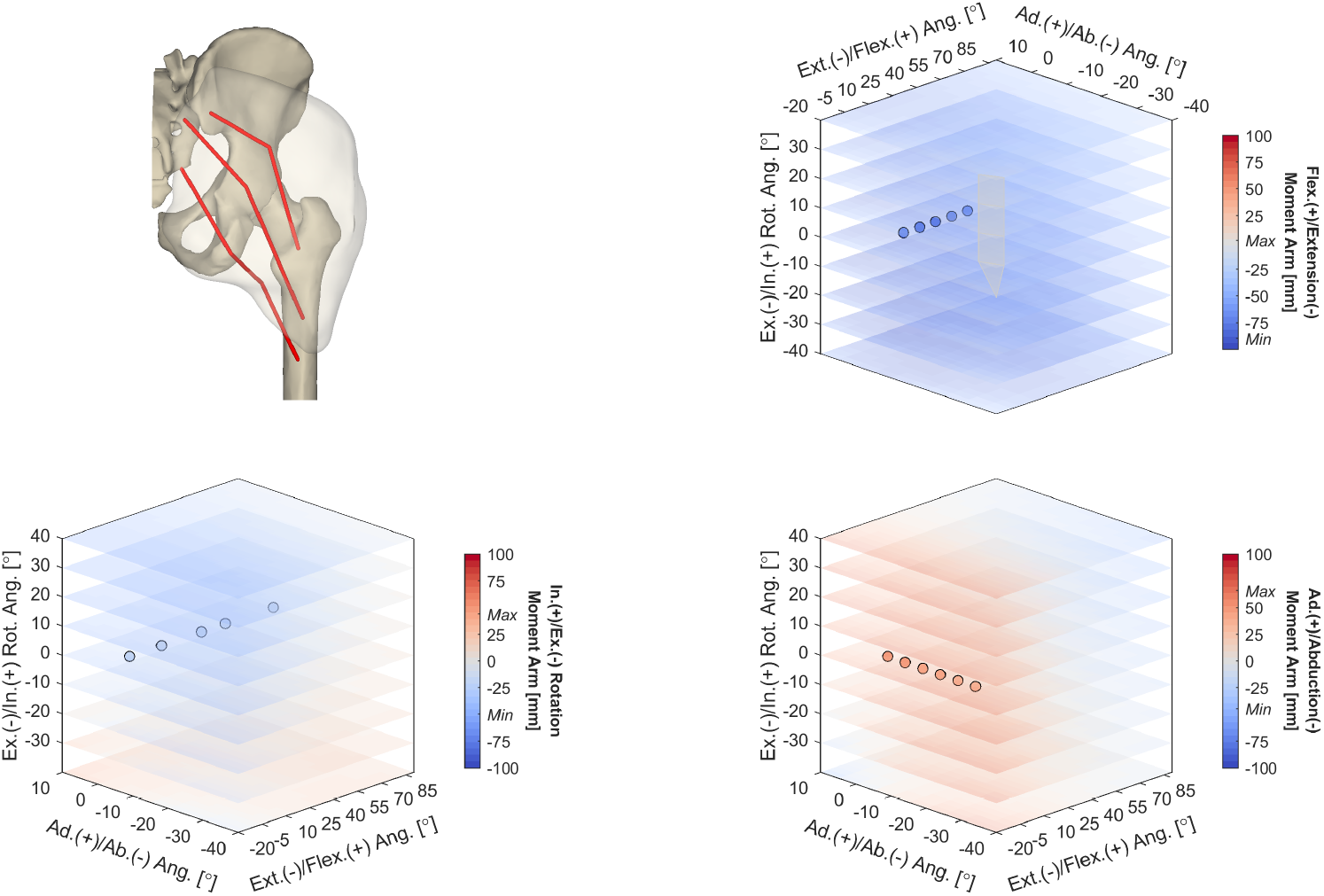
Calibration results of the gluteus maximus. Top left: Paths representing the upper, middle, and lower parts. Top right and Bottom right: 3-D relations of hip extension/flexion and ab-/adduction moment arms with hip angles (lower path); in comparison with measurements of *String 4* from Eng et al. (2015), depicted by the colored scatters. Bottom left: 3-D relation of hip ex-/internal rotation moment arm with hip angles (lower path); in comparison with measurements of *String 6* from Delp et al. (1999), depicted by the colored scatters. The moment arm value is indicated by the color. The gray surface depicts the joint configurations where hip extension/flexion moment arm is greater than −5 mm.

### 4.3. Muscle with No Moment Arm Data

In the complete absence of moment arm data, the calibration must reply on the surface mesh and the designated muscle functions. If a muscle is simple (e.g., the gemelli) and the interested RoM is not too large, the moment arms are practically “clamped” in the correct signs, once the path is configured to go through the muscle mesh. That is, the muscle mesh provides enough information to determine the appropriate functions—notice this is partially the rationale behind the cause-oriented approach of path modeling—and the sign-based cost term could be somewhat redundant.

Nevertheless, such mesh-based “clamping” is often not reliable. If a muscle is large, there will be multiple possibilities of path alignment, some of which could lead to abnormal moment arms. There is a similar issue if the interested RoM covers distant area in the joint space, such as when the joint is posed in extreme positions. Also, a muscle may actuate a motion in both directions, and it will be difficult to tune the path for its moment arm sign to “flip” at certain joint configurations. In these cases, similar to the example of the gluteus maximus, it is necessary to establish additional rules for the moment arm signs.

For example, according to Gilroy et al. (2020), the adductor longus functions as a hip flexor when hip flexion is below 70^°^, otherwise it functions as an extensor. With this, we set it as an objective for moment arm calibration, and as shown in Fig. 9 (top right, gray surface), the function of the adductor longus switches from hip flexion to extension when hip flexion is around 70^°^. Furthermore, considering the major function of the addutor longus, another objective is that it remains a hip adductor throughout the joint space, and the result is shown in Fig. 9 (bottom right).

**Figure 9.**
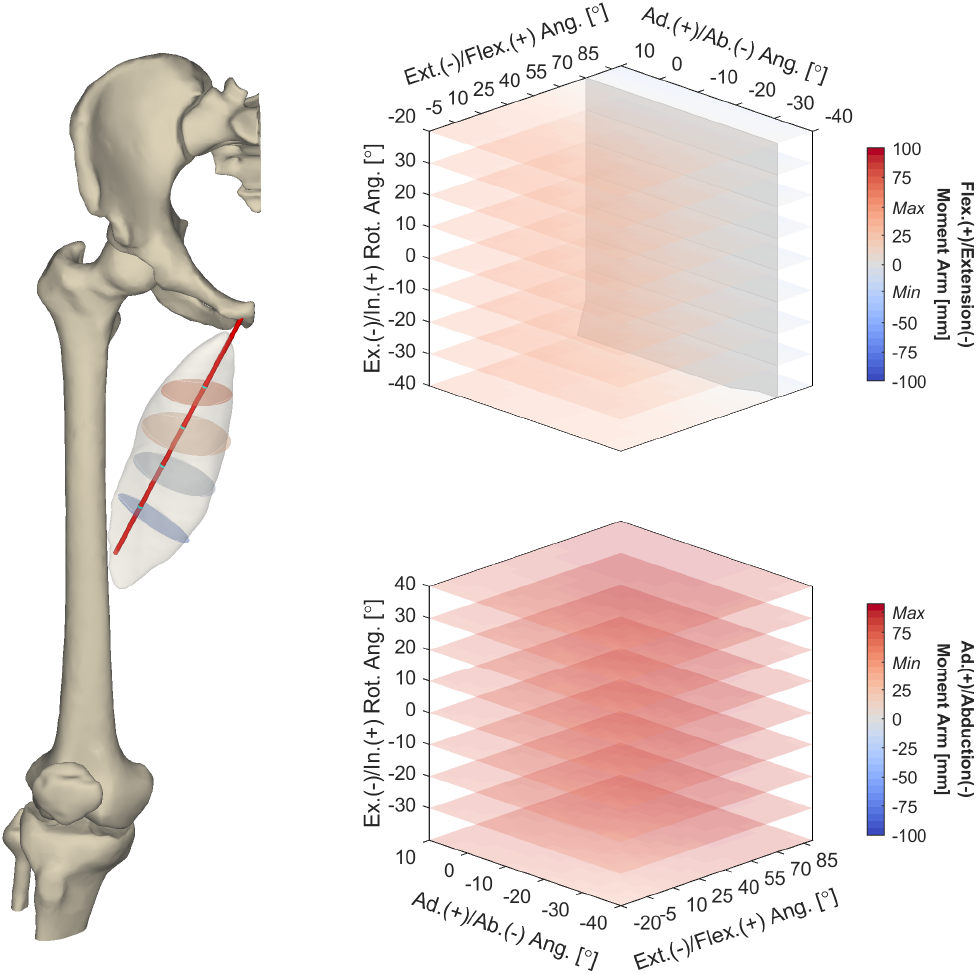
Calibration results of the adductor longus. Left: Ellipse–path intersections (denoted by the cyan nodes). Top right: 3-D relation of hip extension/flexion moment arm with hip angles. Bottom right: 3-D relation of hip adduction moment arm with hip angles. The moment arm value is indicated by the color. The gray surface depicts the joint configurations where hip extension/flexion moment arm is zero.

### 4.4. Joint Kinematics

The muscle moment arm ***r***(***q, p***) depends not only on the path configuration ***p***, but also on the joint kinematics ***q***. Hence, before moment arm calibration is possible, the joint kinematics must be sufficiently accurate. More specifically, the rotation center and axis of each modeled motion should match those of the *actual* motion.

In this work, we transplanted the joint kinematics from the Arnold model (Arnold et al., 2010) to the VHM skeleton (Andreassen et al., 2023): The rotation centers were manually determined by matching the relative positions in the bones, and the rotation axes were replicated. This approach works better for the hip, since the acetabulofemoral joint is a simple ball-and-socket joint. For the knee and the ankle, there are no specific bony landmarks that the rotations center around, and the axes may also differ across individuals. If the motions (or poses) are sufficiently recorded, the modeling of the joint kinematics is not particularly prone to error. Unfortunately, the images for the VHM were obtained in only one pose (Ackerman, 1998), so we cannot determine if (and how) the transplanted kinematics should be corrected.

For the most part, the rotation axes from the Arnold model need no adjustment for the motions to appear natural and for the bone meshes not to overlap within the RoM. However, there is one exception where we must slightly adjust the rotation axis: Regardless where we place the rotation center, the VHM patella does not slide along the patellar surface of the VHM femur as the knee flexes. When moment arm calibration is based on this inappropriate kinematics, the paths for quadriceps insert not onto the patella, but the adjacent void space. Interestingly, when we adjusted the rotation axis so that the patella slides properly on the femur, the insertion points became anatomically realistic (Fig. 4, bottom left).

When it comes to moment arm calibration, there is one kinematics-related problem that can be easily overlooked: The motions (or poses) used to collect moment arm data might not match those used for kinematic modeling. For instance, depending on how ankle ever-/inversion is defined, the motion under this name may have different rotation axes. In the default OpenSim model Gait2392 (Delp et al., 1990) or the Arnold model, the ever-/inversion axis is defined as some combination of all three cardinal axes according to Inman (1976). Nevertheless, when McCullough et al. (2011) and Hintermann et al. (1994) measured ankle moment arms, they mounted ankle specimens on a device to apply motion along each cardinal axis, which is no longer “natural.” So in these experiments, ever-/inversion is defined as the rotation along the coronal axis and essentially a different motion from that modeled in Gait2392 or the Arnold model. Simply put, the same joint configuration (e.g., 20^°^ inversion) does not lead to the same joint position, and the moment arm will also differ.

If we use the moment arm data from McCullough et al. (2011) and Hintermann et al. (1994) to calibrate a model with a “natural” ever-/inversion, paths such as the extensor hallucis longus may route in an unrealistic fashion—e.g., floating above the foot—to compensate the moment arm deficit caused by the kinematic mismatch. To accommodate these data, we modeled an extra set of ankle rotations, which match those used in the experiments. Figure 5 shows the paths for the long digital extensors and flexors, which run along the foot without much deviation.

### 4.5 Identification of Attachment Sites

For path modeling methods such as Modenese and Kohout (2020) and Wang et al. (2025), a prerequisite for full automation is automatic identification of the points (or areas) of origin and insertion. This would require the morphological information of the tendon, which is not always available since the tendon can be thin and hard to distinguish from the surrounding tissues (Andreassen et al., 2023). More importantly, if the mesh of a long tendon is missing, accurate path generation is no longer possible with Modenese and Kohout (2020) or Wang et al. (2025). With our hybrid calibration method, the absence of tendon mesh is not a serious problem, because path alignment is also guided by the moment arms. Take the example of the long digital extensors and flexors, although the meshes of their long tendons are not reconstructed from 3D MRI, the paths still take the appropriate turns in order to yield the correct moment arms (Fig. 5, bottom).

Nonetheless, the lack of tendon mesh is not without an impact on our method. When a path is designated to go through the mesh of the muscle belly, its origin and insertions points are free to locate in a large space. Even with the objective to match moment arm values and signs, the resultant path may still have abnormal attachments, sometimes away from the bones. To make sure that the path attachments are anatomically reasonable, we made estimations based on the meshes of the muscle belly and the relevant bones (see Appendix A) and used them to bound the optimization. If a tendon is long, the absence of its surface mesh will make the estimation inaccurate, which then needs to be corrected with more accurate estimations for other attachment points. For instance, the co-insertion of the gracilis, the sartorius, and the semitendinosus (i.e., the pes anserinus) can be computed from the origins of the tibial muscles. However, we must admit this may not be applicable for all subjects, and machine learning could be a more reliable approach for automated identification of muscle attachment sites.

Lastly, there is one scenario where a single pair of attachment sites is not sufficient. One may have noticed that although Andreassen et al. (2023) reconstructed the mesh for 38 VHM muscles, the results of 42 paths are shown. This is because we modeled more than one path for broad muscles such as the gluteus maximus (upper, middle, and lower), the gluteus medius (anterior and posterior), and the adductor magnus (deep and superficial). As an effect-oriented approach, the main reason why a muscle should be represented by multiple paths is that its strings may have distinct functions. Yet if there are not enough moment arm measurements to guide the additional paths to anatomically reasonable attachments, separate estimations must be made for these sites. For the gluteus medius, path division is easily achieved by inputting moment arm data from the anterior and posterior strings, respectively. This also applies for the glutues maximus, but the attachments of the upper, middle, and lower paths will not spread out as in Fig. 3 (center) without slightly manipulating the estimations. For the adductor magnus, since no moment arm data is available, the modeling of the superficial and deep paths must rely on different estimations of the insertion points.

### 4.6. Strengths and Limitations

Distinct from recently proposed muscle path modeling methods that are automated (Livet et al., 2022; Lloyd et al., 2021; Modenese and Kohout, 2020; Wang et al., 2025), our method is effect-oriented based on the hybrid calibration of both muscle surface mesh and moment arm. The comparison of these methods is shown in Table 1.

**Table 1.**
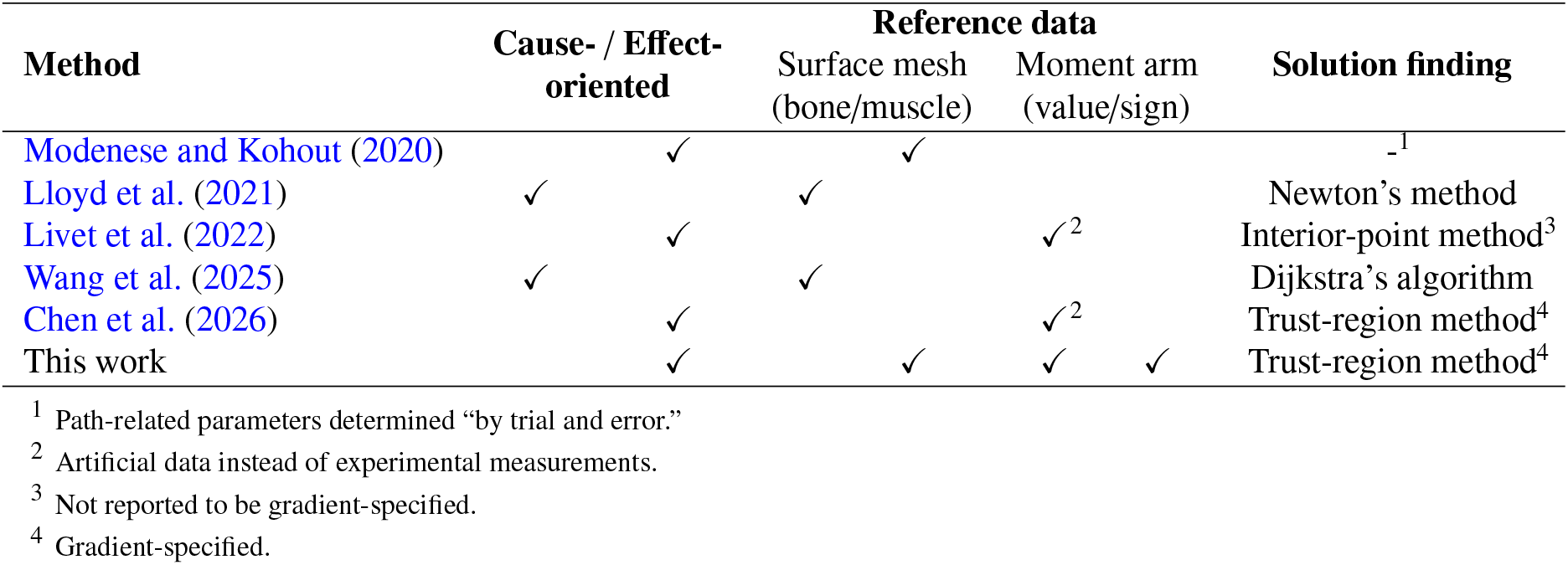
Methods of automated muscle path modeling.

The method of Lloyd et al. (2021) is cause-oriented and is characterized by the direct computation of path wrapping on arbitrary surfaces, which eliminates the need to manually define wrapping obstacles that approximate anatomical obstructions. However, because obstacle contact must be evaluated for a series of points along the path, the method is computationally demanding and may be less practical for large-scale simulations. A simplified cause-oriented approach is proposed by Wang et al. (2025): Instead of computing obstacle contact for a series of points along the path, a set of vertices is identified as wrapping points such that the path length is minimized. In essence, dynamic via points are used to approximate wrapping behavior and thereby reproduce muscle deformation. Nevertheless, the resulting path length varies discretely as wrapping points switch between mesh vertices, which prevents the derivation of an analytical expression for the moment arm.

The method of Modenese and Kohout (2020) may be regarded as effect-oriented in the sense that the generated path is expected to reproduce the muscle length, since the path is posed consistently with the musculotendon mesh. However, the calibration explicitly target the moment arm–joint angle relation, and empirical tuning is required for the path to deform reasonably across RoM. Moment arm–based calibration was presented in Livet et al. (2022). Their optimization framework is not gradient-specified, which limits computational efficiency, and the performance was demonstrated with artificial datasets and relatively simple via point–based paths. It therefore remains unclear how their framework would perform for more complex path structures involving wrapping obstacles, as demonstrated in Chen et al. (2026), or when the calibration data are less comprehensive, as addressed in the present work.

A key advantage of our method is the use of gradient-specified optimization, which improves calibration speed and accuracy compared with using the default numerical gradient (Chen et al., 2026). In a 6-DoF lower-limb model, the calibration of 42 paths took only 20.1 min, and such convenience may encourage model recalibration (or individualization) whenever the measurements are updated. As for calibration accuracy, the final cost for 30 paths are below 0.03 (and 37 are below 0.1), which suggests a good achievement of our threefold objective; see Appendix D for the interpretation of the cost. With manual tuning, it is difficult to match moment arm–joint angle relations as our automated method did. Especially for muscles actuating multiple DoFs, the moment arm relations are high-dimensional, meaning that they are difficult to visualize, let alone manually calibrate.

In the spirit of automation, we made an effort to remove as much manual work as possible, including the estimation of the origin and insertion points. As we previously emphasized, due to the lack of tendon surface mesh, it could be difficult to make accurate estimations for certain attachment points using the method in Appendix A. Despite the possibilities to correct the estimations automatically, the solutions are format-wise less neat and the applicability is uncertain due to subject variability. In the future, we look to develop automatic identification of the origin and insertion points (or areas) with machine learning. This would not only simplify the preparation step in our method, but also make the methods of Modenese and Kohout (2020) and Wang et al. (2025) fully automatic.

Apart from the morphological data, there are also issues related to the moment arm data. First, our model is based on the musculoskeletal geometry from the VHM, but the moment arm data for calibration come from other studies—the difference in the subject group might induce errors for our hybrid calibration. Second, the moment arm data may also come from different experiments, each with its own subject group, method for measurement, and nuance in the definition of joint rotations. At last, many muscles have limited or even no data for calibration. These data-related issues are currently inevitable, and the paths in this study will need to be validated or recalibrated, if more comprehensive datasets are available in the future.

On the topic of validation, this is unfortunately absent in our work. Ideally, validation of the paths would need muscle surface meshes or moment arms in additional joint configurations, but it can be technically challenging to collect either type of data across a large RoM. However, if we are to envision, the more practical goal would be to focus on the morphological data. Our modeling results of musculoskeletal geometry (Figs. 3–5) are based on images obtained in only one pose. If the meshes from a series of diverse poses are available, then the path may be configured to go through multiple sets of cross-sectional ellipses. The geometric features of the muscle will naturally be better reflected in a wider RoM. More importantly, the moment arms of such a path are somewhat reliable even without calibration. The rationale is that, by going through the muscle mesh in each joint configuration, the length of the path changes in a similar fashion as that of the muscle does. In other words, the derivative of muscle length with respective to joint angle (namely moment arm) is similar, and this is essentially the mechanism of the *tendon excursion* method for moment arm measurement (An et al., 1984; Maganaris et al., 2000).

## 5. Conclusion

A gradient-based optimization method is proposed for automated muscle path modeling and tested with experimental data from literature. As an effect-oriented approach, this method is based on the hybrid calibration of muscle surface mesh and moment arm, and the resultant paths are both anatomically realistic and biomechanically accurate. With its speed and accuracy, our automated framework is suitable for large-scale subject-specific applications.

## 6. Acknowledgements

This work was supported by the Lighthouse Initiative Geriatronics by StMWi Bayern (Project X, Grant No. 5140951).

## Appendix A. Identification of Entry and Exit Points

Given sufficient knowledge in the human anatomy, it is not difficult to manually identify the origin and insertion areas on the bone meshes or the areas on the muscle mesh adjacent to the origin and insertion. However, in preparation for subject-specific applications at large scale, our concept of automation is to remove as much as manual work as possible. Below we propose an algorithm for automatic identification of entry and exit points, and the results based on Andreassen et al. (2023) are generally satisfactory, despite the lack of tendon mesh.

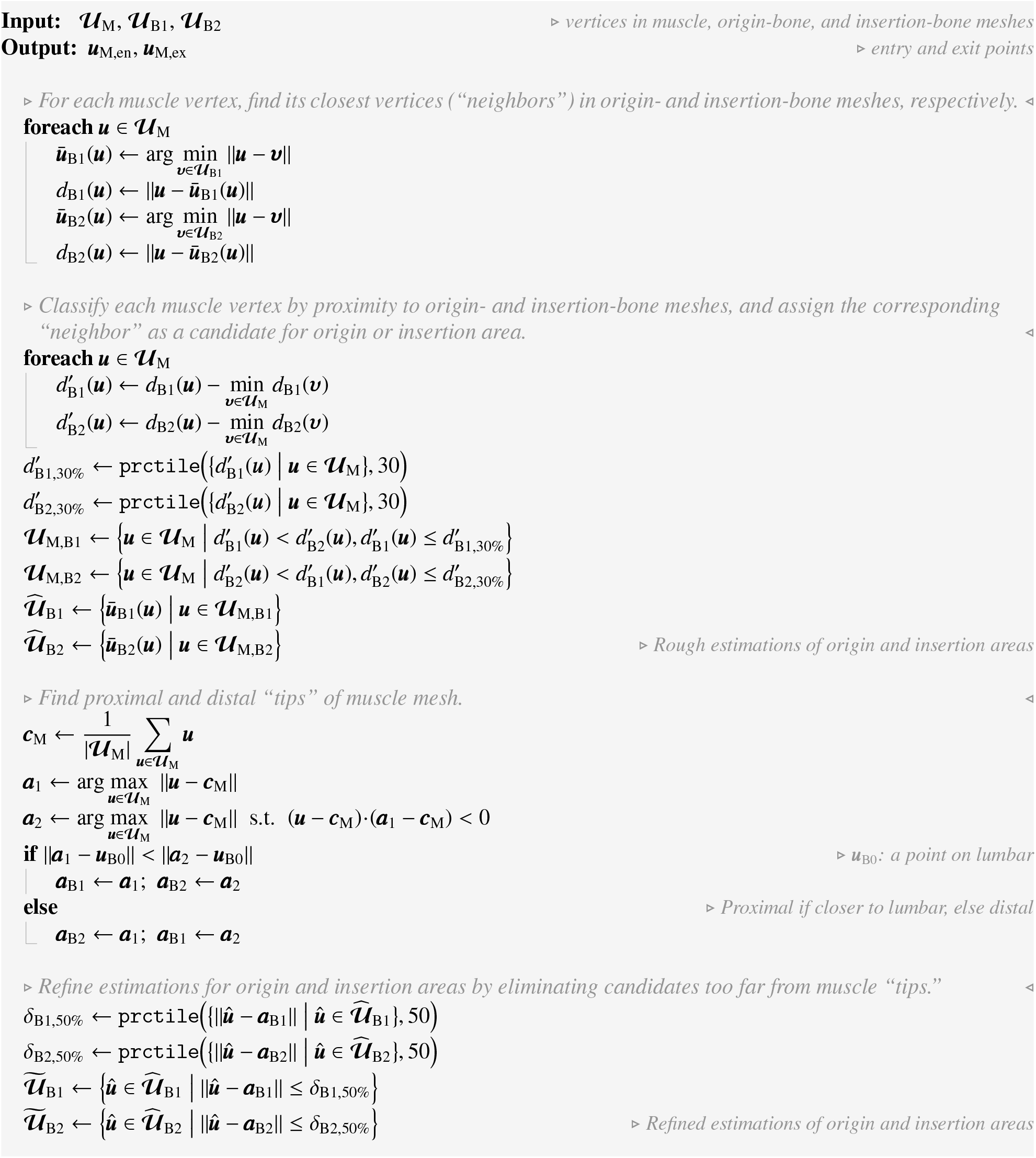
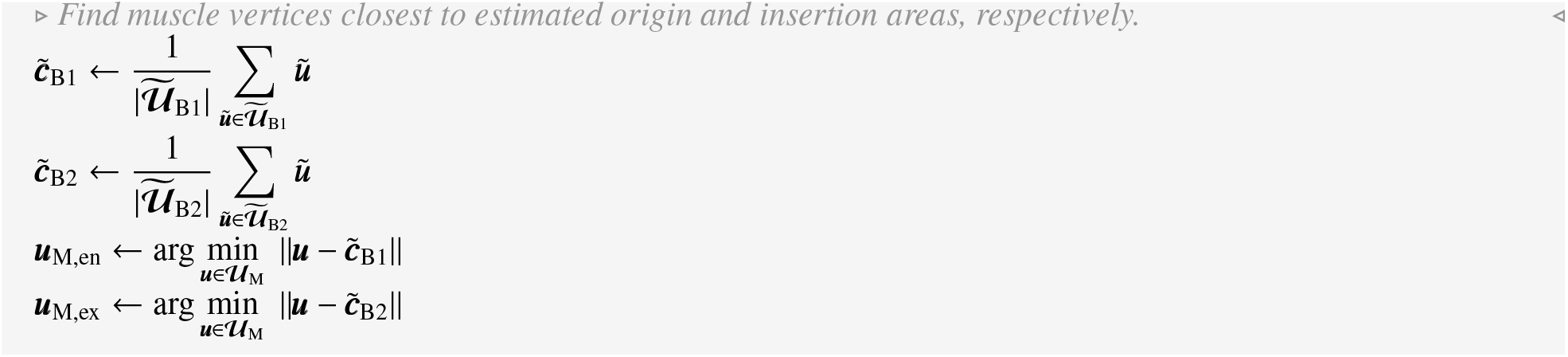

Here, 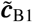and 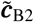 may also serve as an initial guess of the origin and insertion points. The estimation is particularly accurate if the tendon is short or if its mesh is available. However, if the mesh of a long tendon is absent, 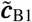 or 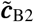 might deviate from the correspondent anatomical landmarks. In this case, there are various ways to correct them automatically, but we will not elaborate on the details here. Essentially, as a by-product for the entry and exit points, we estimate the origin and insertion points. Notice they are compatible with the path modeling method of Wang et al. (2025), whose inputs of “attachment points” are not obtained automatically. Similarly, in Modenese and Kohout (2020), the inputs of “attachment areas” are not reported to be obtained automatically, for which 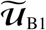 and 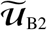 serve as a good estimation.

## Appendix B. Ellipses for Muscle Cross-Sections

With the entry and exit points (***u***_M,en_ and ***u***_M,ex_ in Appendix A) as boundary constraints, we apply the method of Dong et al. (2005) to construct a harmonic scalar field *h*(·) over the muscle mesh. Depending on the relative position between the entry and exit points, each of the *N*_M_ vertices is assigned a scalar value *h*(***u***_M,n_) ∈ [0, 1], forming a series of field isolines. Similar to Modenese and Kohout (2020), we treat these isolines as the contours of muscle cross-sections, but we do not thread the path through any certain location (e.g., the geometric center) on multiple cross-sections. Instead, we only expect them to be intersected by the path, so that the path has enough room to be tuned for moment arm calibration.

For computational convenience, we simplify each cross-section as an ellipse, which is geometrically similar but computationally easier to implement. To obtain these cross-sectional ellipses, we first extract *N*_ell_ vertex subsets, each from a narrow belt formed in-between a pair of isolines.

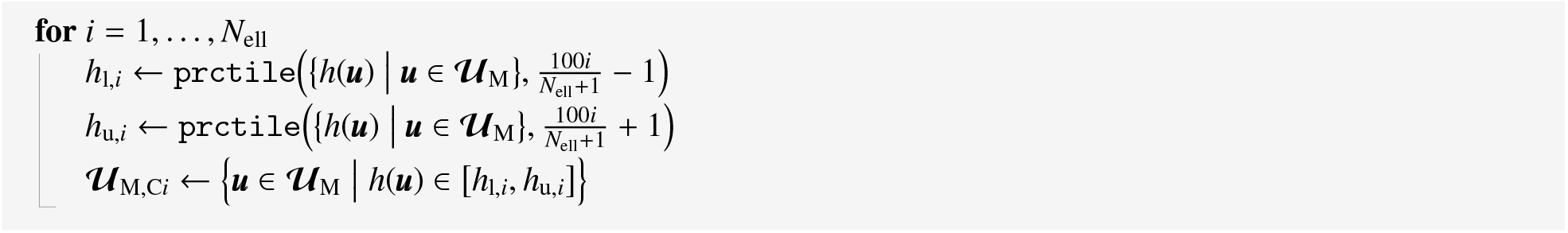

Then, the 3-D points in each subset are projected onto a 2-D plane, whose coordinates are denoted by ^ϵ^ (·), using singular value decomposition, and the 2-D points are fitted with an ellipse with the method of Fitzgibbon et al. (1999).

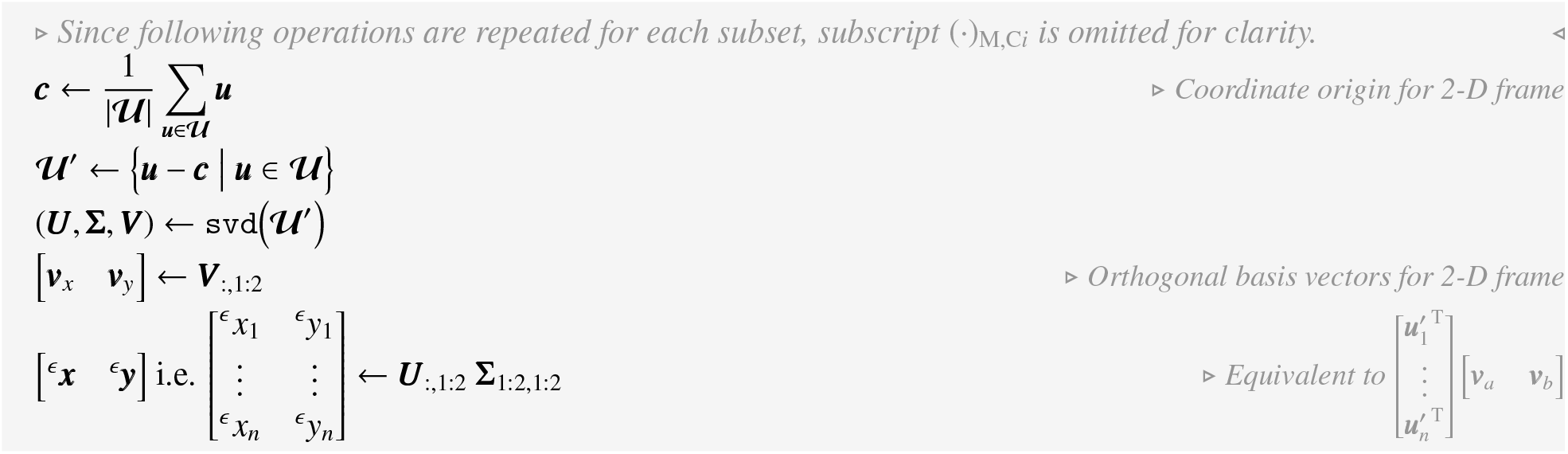
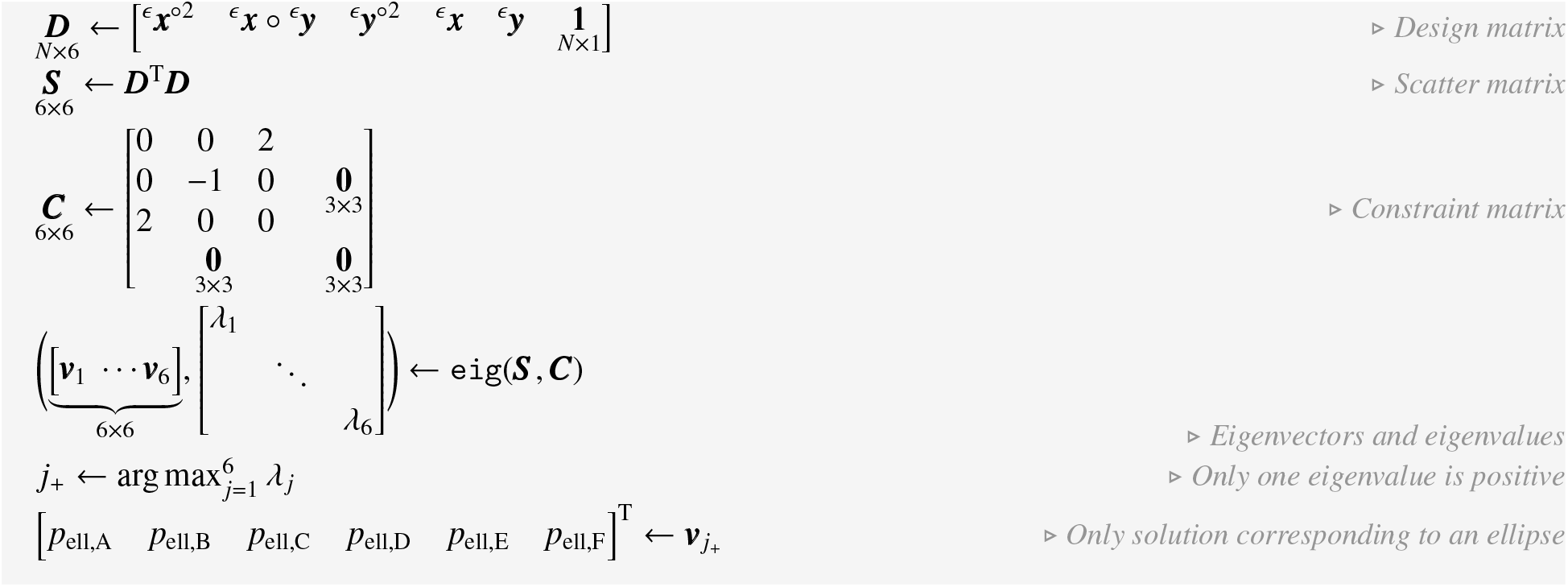

Here, the solution is a 6-D vector, whose elements correspond to the six conic coefficients defining an ellipse. To improve readability in the main text, we convert this general form into a canonical form. More specifically, the coordinate frame is translated so that its origin coincides with the ellipse center and rotated so that its *x*- and *y*-axes align with the ellipse’s major and minor axes, denoted by ^e^(·).

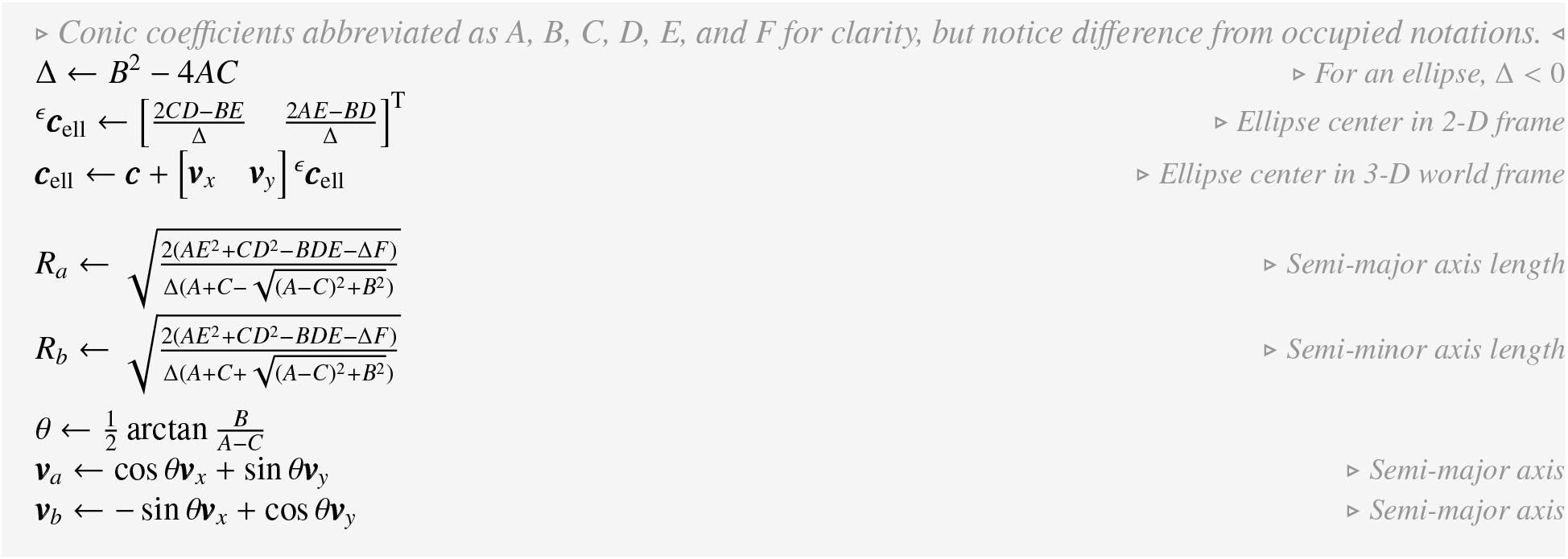

General form:

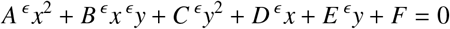

Canonical form:

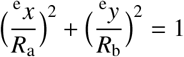

Transformation into 3-D world frame:

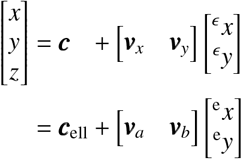

## Appendix C. Ellipse–Sub-Segment Intersection

For each ellipse representing a muscle cross-section, it is expected to be intersected by at least one sub-segment of the path. Depending on both the path structure and the joint configurations, a path can be divided into one, two, three, or six sub-segments. A sub-segment is straight if no wrapping occurs between two anchor points, otherwise it is curved. However, when the intersection is computed, we treat all curved sub-segments as straight. This significantly simplifies the computation, and since none of the lower-limb muscles are heavily wrapped in the fixation position, the difference will be small.

Suppose ***u***_AP1_ and ***u***_AP2_ are the two anchor points constituting a straight sub-segment, the intersection with the ellipse must be along the sub-segment “ray”

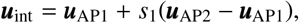

where *s*_1_ is a scalar indicating the location of ***u***_int_ relative to the sub-segment. As for the ellipse, it lies in a plane spanned by ***v***_*a*_ and ***v***_*b*_, and the intersection point must also be on this plane and satisfy

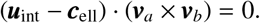

First, we substitute one equation into the other to have

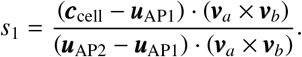

If 0 < *s*_1_ < 1, the intersection takes places between the two anchor points, otherwise the plane can only be intersected if the sub-segment is extended along either side.

Then, we compute a second indicator, which describes the location of ***u***_int_ relative to the ellipse

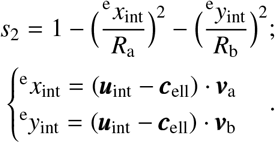

If *s*_2_ > 0, the intersection takes places within the ellipse, otherwise the sub-segment ray can only be intersected if the ellipse is expanded across the plane.

## Appendix D. Cost Normalization

The optimization problem we formulate for muscle path modeling is multi-term objective: Each of the three objectives targets at a different property, and the cost terms need to be normalized so that their contributions can be meaningfully combined.

The first key of normalization is to ensure that the cost terms have the same units. In Eq. 2, the cost term for muscle surface mesh *J*_SM_ is based on the quantity of intersected ellipses, which is dimensionless. When matching moment arm values, *J*_vMA_ in Eq. 7 is based on moment arm length, with the unit of mm. For this, the difference of model and measured moment arms is divided by a tolerance value, also with the unit of mm—this ratio is dimensionless. When matching moment arm signs, *J*_sMA_ in Eq. 8 is based on the quantity of matched signs, which is already dimensionless and does not require normalization.

Then, the magnitude of each cost term needs to be scaled as an approach to balance the importance of each objective. In Eq. 2, we assign a factor of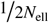 so that the maximum of *J*_SM_ is 1/2 when none of the *N*_ell_ ellipses are intersected. This sets a baseline for the “worst-case scenario” in surface mesh calibration, and the other two cost terms may be weighted with respect to this baseline. In this study, we intend to treat surface mesh and moment arm as equally important. In other words, the sum of *J*_vMA_ and *J*_sMA_ should be scaled to 1/2 when the value-based calibration and the sign-based calibration are both in their “worst-case scenarios.”

In the sign-based moment arm calibration, the “worst-case scenario” is simply when none of the signs match, and the cost term will reach its maximum

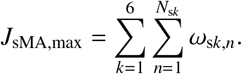

For the value-based moment arm calibration, the situation is more complicated. Since there is no limit to the difference between *r*_target_ and *r*_model_, the theoretical “worst-case scenario” will drive *J*_vMA_ to infinity. Hence, a nominal “worst-case scenario” is needed, and we define it as when the moment arm error reaches the designated tolerance of (10% |*r*_target_ |+ 5) mm; accordingly,

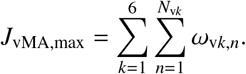

Suppose the datasets for moment arm calibration are from *N*_ref_ different sources. For the *j*-th dataset, it contains *N*_d *j*_ data, each of which is a target value or sign for one of the six potential moment arms at a certain joint configuration. From another perspective, for the *k*-th moment arm, it needs to match *N*_v*k*_ target values and *N*_s*k*_ target signs at the designated joint configurations. Thereby, there is a one-to-one correspondence between the associated weights of each datum in a dataset (ω_d *j,m*_) and each target for a moment arm 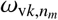 or 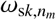. Then, in the “worst-case scenarios,”

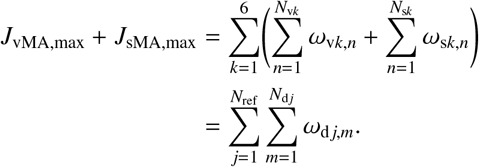

For this sum to match the baseline of 1 set in the surface mesh calibration, simply have

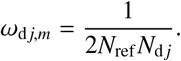

This is equivalent to weighting all data within each dataset equally, while weighting all reference sources equally regardless of the dataset size.

With this normalization, the cost attains a meaningful interpretation that is consistent for any muscle: If *J* = 1, it means that

1. when posed in the fixation position, the path fails to go through any of the ellipses,
2. at the designated joint configurations, the model moment arms deviates from the target values by the designated tolerances, and
3. at the designated joint configurations, the model moment arms are opposite from the target signs.

The calibration results may now be quantitatively compared across different muscles, and we may also set a common cost threshold (e.g., 0.03) to indicate if a muscle path is well-modeled.

## Appendix E. Initialization and Configuration for Optimization

In order for the optimization to proceed sufficiently fast without compromising accuracy, we follow the configuration in our recent work (Chen et al., 2026):

- `trust-region-reflective with SpecifyObjectiveGradient;`
- 10^−4^ for `StepTolerance`, and 2.5 × 10^−16^ for `FunctionTolerance` and `OptimalityTolerance`;
- 10^2^ for `MaxIterations`, and `Inf` for `MaxFunctionEvaluations`.

When ***p*** was initialized for Eq. 1 and later optimized, the locations of the origin and insertion points were bounded within a uniformly distributed range of ±10 mm from 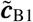 and 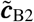 (see Appendix A), respectively. This is because 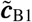and 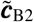 correspond well to the anatomical landmarks of the tendinous attachments, and we include this constraint to speed up the calibration. Importantly, without this constraint, the origin and insertion points might be tuned away from the bones. Although this does not necessarily interfere with moment arm calibration, the path will appear anatomically unrealistic. For other parameters, there are no constraints during the optimization.

Also, when generating the initial points for the path structures of OCI and OC_1_C_2_I, the cylinder centers were randomly initialized around the sections of the line between the initial origin and insertion points. The rationale is to ensure the cylinders are not too departed from the initial muscle paths: If a cylinder is not wrapped at the beginning of the optimization, the relevant gradient will remain a null matrix and the cylinder center might be left untuned till the end. For OVI and OV_1_V_2_I, the via points are initialized in a similar fashion for consistency.

## Appendix F. Data and Results for Moment Arm Calibration

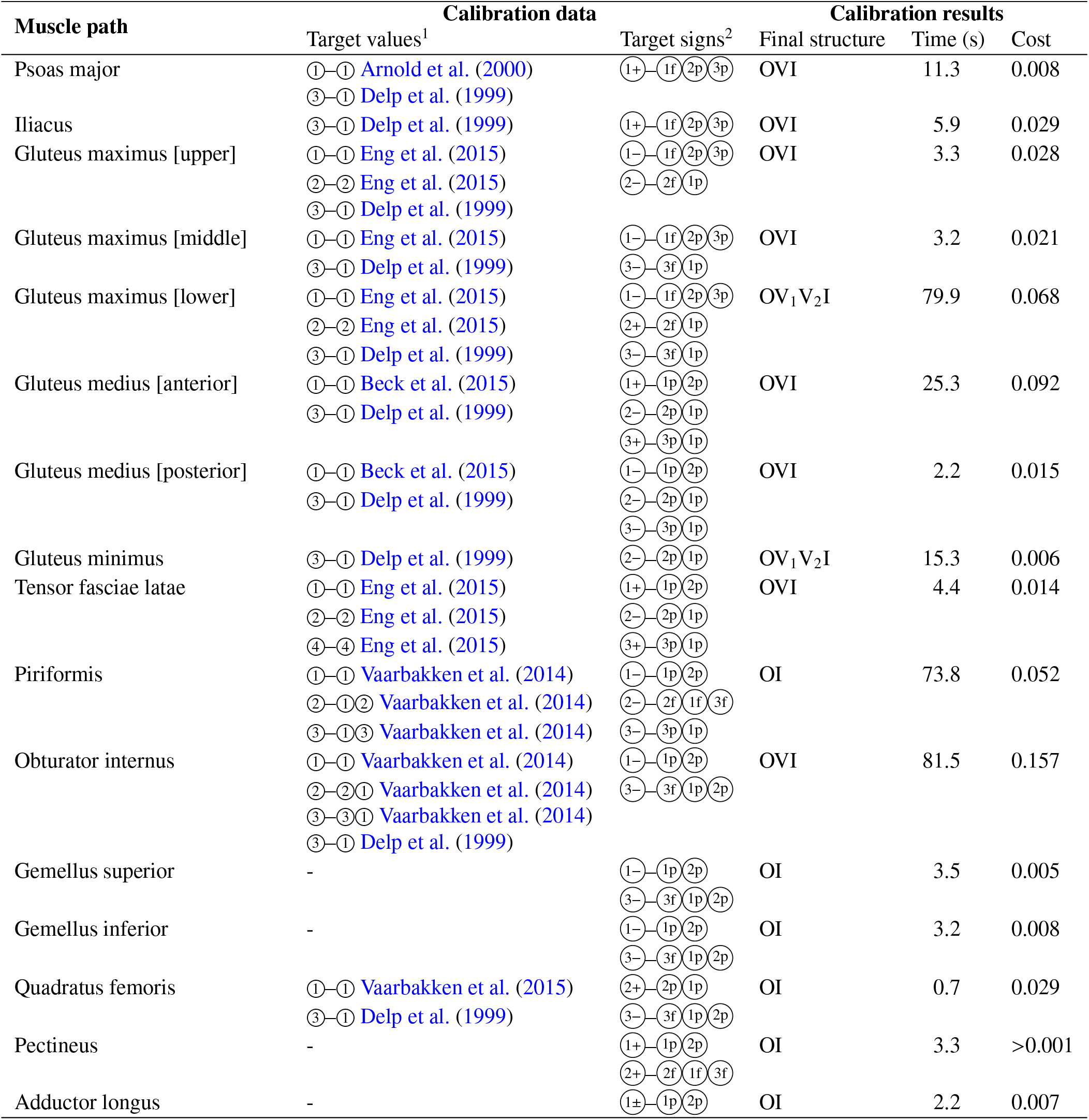
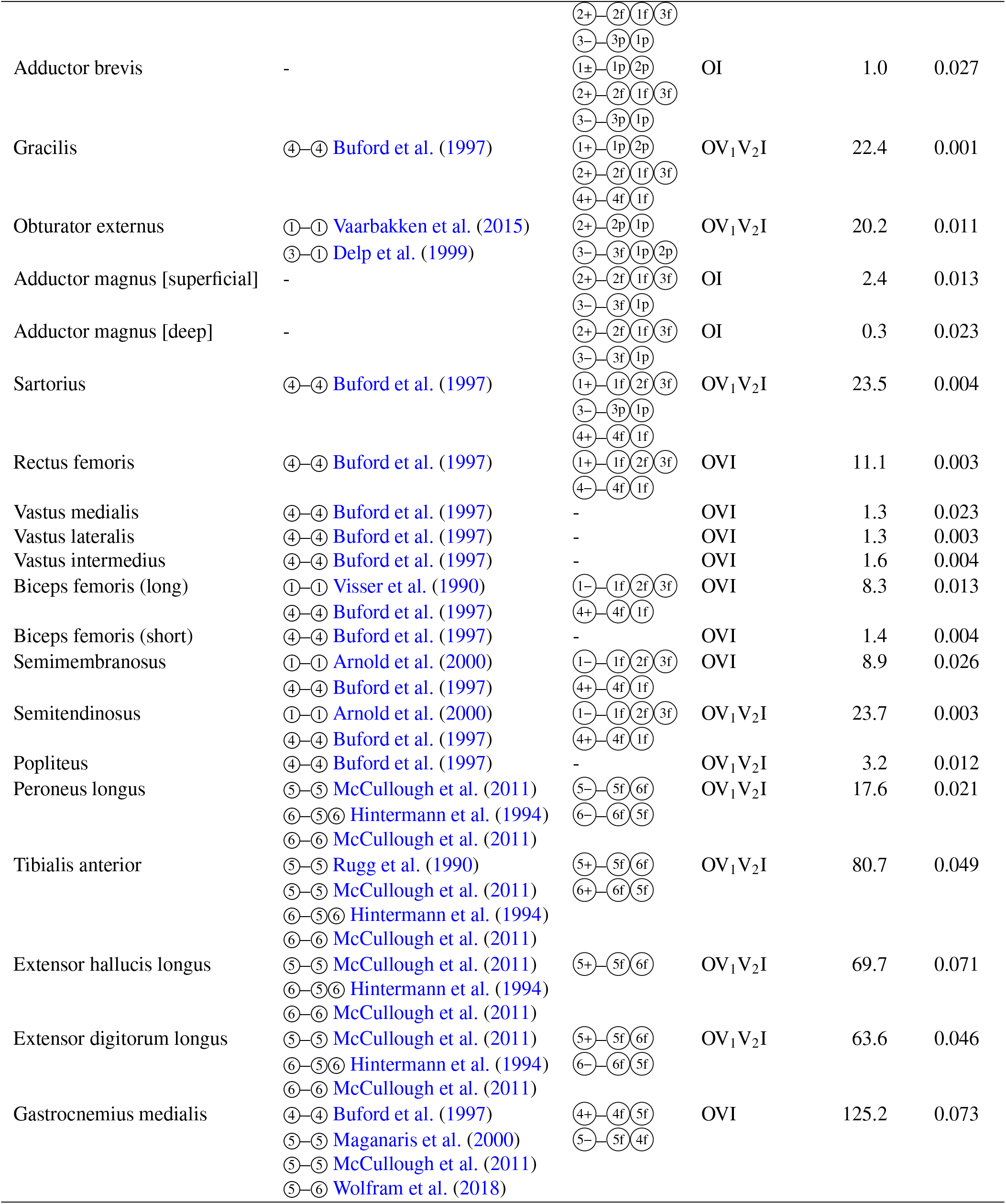
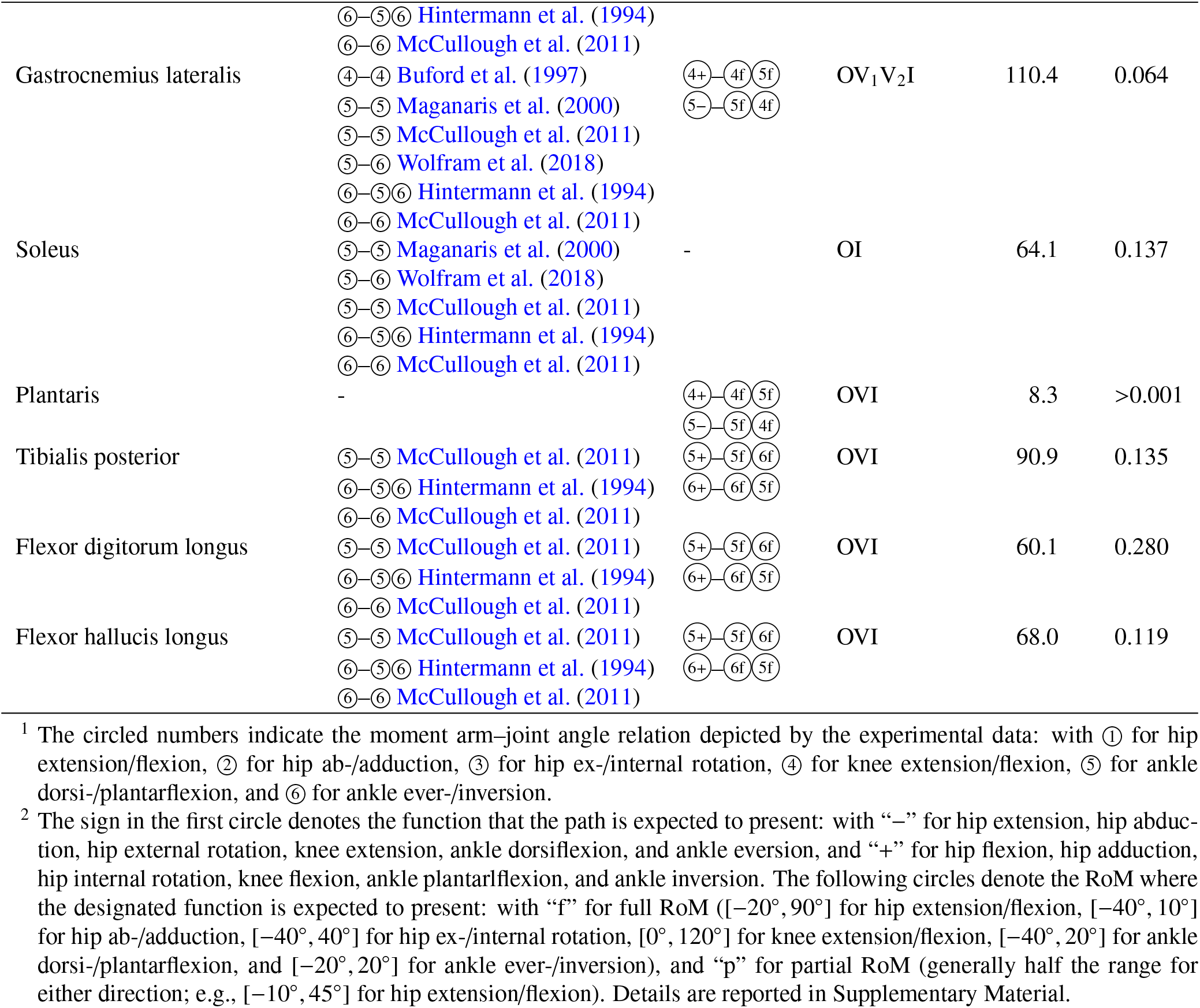

## Footnotes

1 Following the convention in biomechanics, the musculotendon unit is referred to simply as a *muscle*, and the same rule applies when the term is used attributively (e.g., *muscle* length, moment arm, or path). Exception in this article: the geometry (or surface mesh) of the muscle is explicitly distinguished from that of the tendon.

